# Neural control of coordinated wing and leg movements during a terrestrial threat display

**DOI:** 10.1101/2025.10.25.684556

**Authors:** Shuo Cao, David J. Anderson

## Abstract

Threat displays are a common form of social communication. In flying species, such displays often involve stereotypical, choreographed wing and leg movements. How the brain coordinates characteristic movements of these appendages to transmit social signals is poorly understood. In male *Drosophila*, threat displays flexibly combine wing displays with rapid turns and charges. Here we identify two brain modules downstream of a superordinate threat command center. Each contains two neurons that combinatorially generate threat-specific appendicular movements: one comprises two descending neurons synergistically controlling wing threat; the other contains two interneurons that antagonistically control turns and charges. Within-module neuronal coactivation evokes appendage-specific actions recapitulating those evoked by the upstream center. These data uncover a hierarchical combinatorial circuit that coordinates wing and leg movements during a terrestrial social display. More generally, our findings identify an instantiation of a Tinbergenian hierarchical behavioral control system for social communication and reveal a novel underlying implementation logic.

## Introduction

Social displays are communicative behaviors prevalent across the animal kingdom and play a crucial role in encounters such as mating, aggression, and defense^1–9^. A key feature of these displays is that they comprise multiple actions involving the coordinated use of distinct body parts. For instance, during a threat display, elephants spread their ears, raise or swing their trunks, shake or nod their heads, and stomp their feet. The precise coordination of these movements is vital for effective social signaling. While significant progress has been made in identifying neural nodes that control complex social behaviors at a more holistic level^10–15^, much less is known about how individual actions involving distinct appendages are orchestrated to produce complex social behaviors.

This problem is of particular interest in flying species that exhibit terrestrial social displays combining coordinated, stereotyped movements of the wings and legs, appendages which are otherwise utilized independently for aerial and earthbound locomotion. Threat display by *Drosophila* males is an aggressive or agonistic behavior, composed of at least four distinct actions: wing elevation, pump, turn, and charge^16^. Wing elevation comprises an abrupt bilateral upward wing movement to an angle of 45°, which is then sustained for seconds (Figure 1Ai). Pump is a transient (<0.1 s) bilateral horizontal wing extension towards 90° (Figure 1Aii). Turns and charges are both transient and discrete locomotor events that are performed as the fly rapidly orients and accelerates toward its opponent (Figure 1Aiii and 1Aiv). These actions can occur in various combinations: stable wing elevation can be performed with or without turns and/or charges, while a pump is always accompanied by turning and charging^16^. Together these movements visually communicate a larger size and aggressive intent to a male conspecific. This behavioral complexity necessitates a sophisticated neural control mechanism to flexibly utilize and coordinate wing and leg movements during such threat displays.

**Figure 1.**
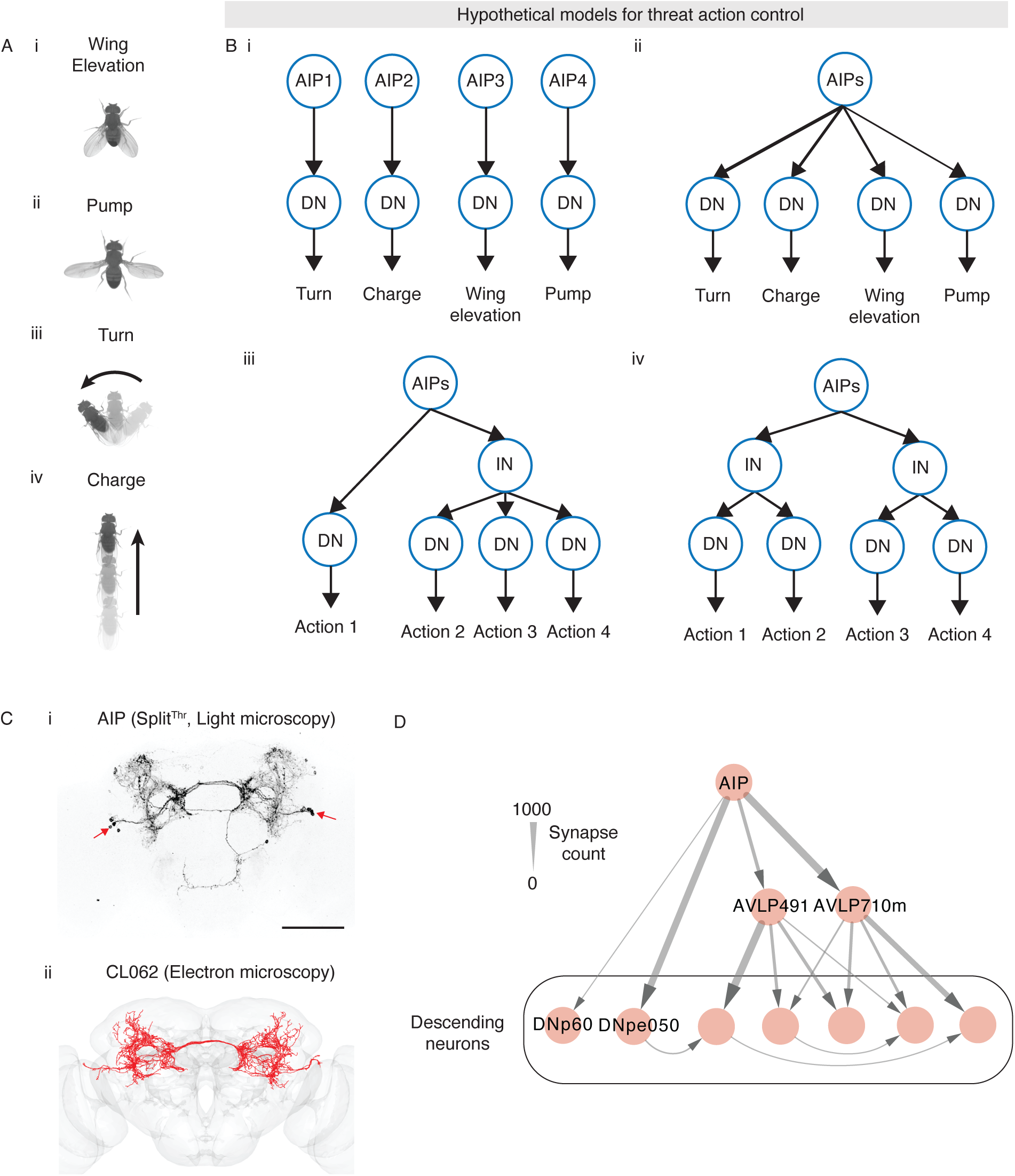
AIP downstream circuit and models of how threat actions are controlled. (A) Four distinct actions of threat display: wing elevation (i), pump (ii), turn (iii), and charge (iv). (B) Models of how AIP neurons control four distinct threat actions. In (ii), thickness of arrows is proportional to strength of connections. (C) Expression pattern of Split^Thr^ (i), and cell skeletons of AIP neurons reconstructed in the male CNS connectome (ii). Scale bar, 100μm. Arrows indicate the cell body locations of AIP. (D) A circuit diagram of AIP and major downstream descending neuron (DN) and interneuron (IN) targets. IN targets that do not project primarily to DNs are omitted for clarity. See also Figure S1.

Previously, we identified a small cluster of genetically labeled interneurons (INs) in the central brain, termed “AIP”, using an unbiased functional activation screen for aggression-promoting neurons^15–17^. Unlike other aggression-promoting neurons identified to date^15,18–25^, the AIP cluster specifically controls only wing threat behavior. Silencing AIP neurons suppresses natural wing threat displays without impairing contact-mediated aggressive behaviors such as lunging, tussling, or boxing^16,26^. Conversely, optogenetic activation of AIP neurons elicits the full complement of threat actions in solitary flies^16^. These data identify AIP as a crucial command center for coordinating threat displays. However, how the activity of AIP neurons is flexibly routed to orchestrate the distinct behavioral actions that comprise these displays remains unknown.

To address this question, we have investigated the logic of threat circuit organization and function downstream of the AIP cluster. Using EM connectomics and functional imaging, we reveal that AIP neurons provide both direct and indirect feedforward input to distinct sets of descending neurons (DNs) controlling the wings or legs, respectively. Through optogenetic and behavioral analysis with cellular epistasis tests, we show that the direct pathway comprises two DNs that synergistically control wing elevation and pump. In contrast, the indirect pathway comprises two functionally antagonistic INs, whose conjoint activity controls a different group of DNs that mediate leg-based threat actions. Together, these findings reveal a hierarchical and combinatorial circuit logic for the coordination of wing and leg movements during a terrestrial social display.

## Results

### Identifying the AIP downstream circuit in a male central nervous system (CNS) connectome

Since AIP neurons are INs located in the central brain, they likely promote wing threat actions by engaging downstream circuits that channel AIP activity into distinct motor pathways, ultimately via the activation of various types of DNs. *A priori* several models could explain the coordination of different appendicular movements. At one extreme, the AIP cluster could be comprised of different neuronal subtypes, each of which controls a distinct motor action via distinct DNs (Figure 1Bi). Such a model would provide maximal flexibility in orchestrating different combinations of threat actions. At the opposite extreme, each AIP neuron could contribute to the control of all four threat actions (Figure 1Bii), perhaps generating flexible action combinations via connections of different synaptic strengths to different DNs. Between these extremes, various types of hierarchical models are possible (Figure 1Biii-iv).

We mapped AIP neurons, labeled by a specific split-GAL4 driver line^16^ (Figure 1Ci), into a male CNS connectome^27,28^ (https://neuprint.janelia.org/, male-cns:v0.9) and identified five neurons per hemisphere with morphological resemblance to the genetically labeled AIP neurons (CL062, Figure 1Cii). We then mapped the major direct and indirect downstream targets of these five neurons, constructing a circuit diagram that incorporates these major targets (Figure 1D and S1A). The resulting diagram immediately excluded the two extreme models, in which either different AIP neurons each control one of the four distinct threat motor actions by projecting to different DNs (Figure 1Bi), or in which all AIP neurons fan out to directly engage a common set of DNs in parallel (Figure 1Bii). Instead, we found that AIP neurons as a group project directly to DNs (DNpe050 and DNp60), as well as to INs (AVLP491 and AVLP710m) which in turn project to various DNs (Figure 1D). The overall AIP-DN network architecture is feedforward. Other IN targets of AIP do not project primarily to DNs (Figure S1A). While further analysis revealed two subsets of AIP neurons (Figure S1A), those subsets projected to highly overlapping downstream targets, differing primarily in the strength of those connections (Figure S1B).

These connectomic data suggested various types of hierarchical models for the control of threat actions by AIP neurons (Figure 1Biii and 1Biv), where some or all threat-related DNs are organized by INs to control subsets of actions. To differentiate among these and other models, it was necessary to gain genetic access to individual AIP downstream targets. Using NeuronBridge^29–34^, we screened a total of ∼70 split-GAL4 lines (Figure S1C) and were able to identify specific drivers for DNpe050, DNp60, AVLP491 and AVLP710m (Figure 1D). As described below, all of these drivers labeled cells that exhibited behavioral phenotypes in silencing or activation experiments and/or calcium responses to AIP stimulation. Although AIP neurons also project to some other classes of INs or DNs (Figure S1A and S1B), we excluded those from further analysis either because we were unable to identify clean genetic drivers for those cells, or because the neurons (VES041^35^) did not reveal calcium responses to AIP activation, or functional effects on threat behavior.

### DNpe050 activation increases the probability of wing elevation

DNpe050 is a DN that receives the strongest monosynaptic input from AIP among all identified DN targets (Figure S1A, S1B, 2Ai and 2Aii) and is predicted^36^ to be cholinergic based on the connectome. An initial round of screening identified a split-GAL4 that labeled DNpe050 (VT058429-AD, 12G09-DBD), along with approximately 40 additional neurons per hemisphere. To reduce off-target labeling, we performed enhancer bashing^22,37,38^ on both hemi-drivers (Figure S2A) and tested 10 different combinations of enhancer fragments and one of the parental drivers (Figure S1C, DNpe050 EB drivers). Two of these newly created hemi-drivers, VT058429EB1-AD and 12G09EB3-DBD, retained expression in DNpe050 when combined, labeling only ∼10-15 additional neurons. Further screening using these enhancer-bashed hemi-drivers and additional hemi-drivers from NeuronBridge (Figures S1D, DNpe050 second round) yielded several split-GAL4 combinations of which four, including DNpe050-split1 and DNpe050-split2, were further characterized. When triply intersected with Otd-FLP^21^ to eliminate ventral nerve cord (VNC) expression, DNpe050-split1 labeled only 2-3 off-target neurons (Figure 2B) while DNpe050-split2 labeled a group of approximately 5-8 off-target neurons distinct from those labeled by DNpe050-split1 (Figure S2B).

**Figure 2.**
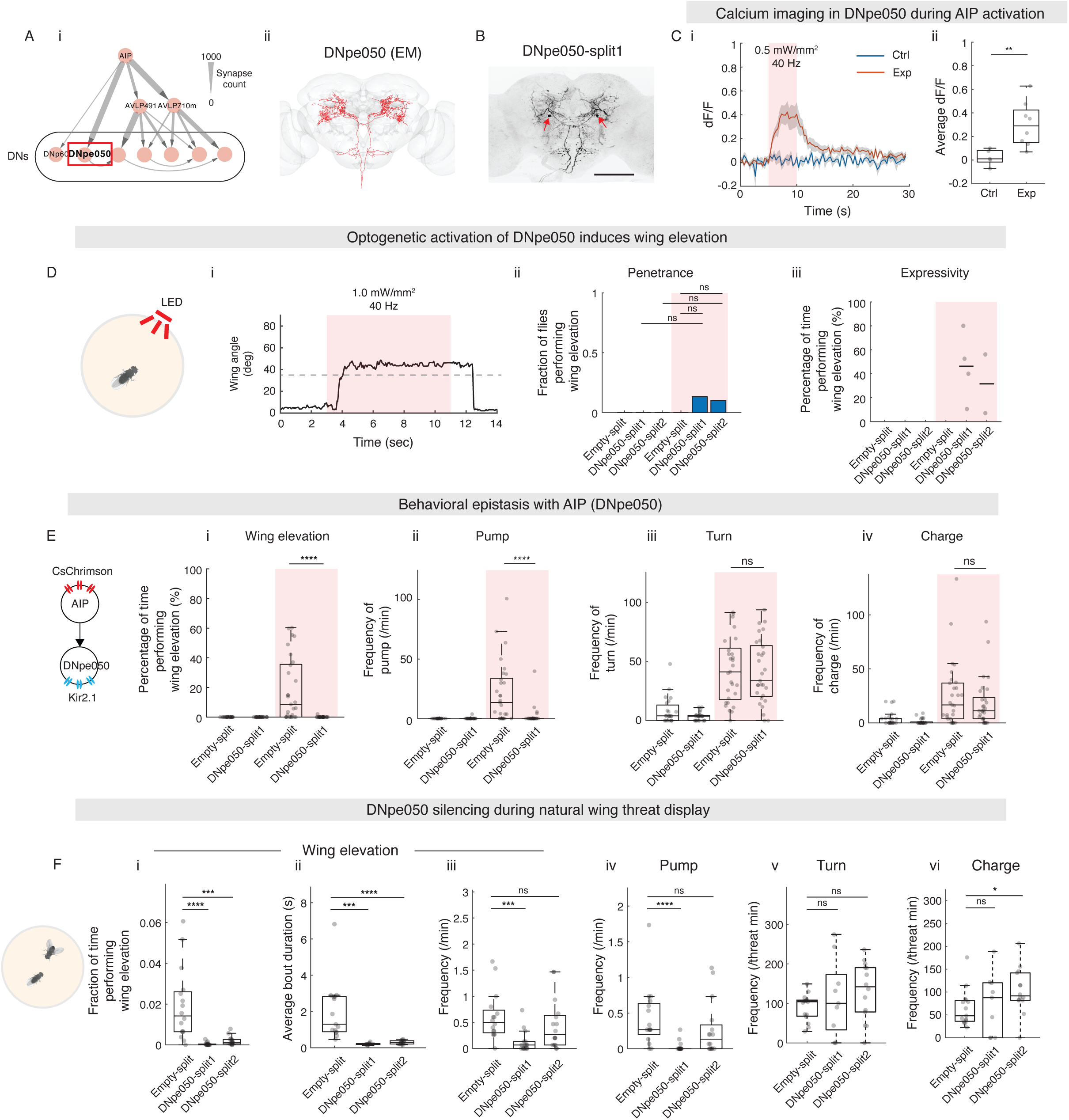
DNpe050 activation increases the probability of wing elevation. (A) (i) DNpe050 highlighted in the AIP downstream circuit diagram (red box). (ii) Cell skeletons of DNpe050 reconstructed in the male CNS connectome. (B) Expression pattern of DNpe050-split1 (with Otd-FLP). Arrows: cell body locations of DNpe050. Scale bar, 100μm. (C) jGCaMP8m fluorescence changes in DNpe050 in response to AIP stimulation in head-fixed living flies. (i) DNpe050 response (dF/F) before, during, and after photostimulation of AIP. Red shaded region indicates light delivery. (ii) Average DNpe050 response (dF/F) during photostimulation. Exp: AIP > Chrimson; DNpe050-split2 > GCaMP8m. Ctrl: Empty > Chrimson; DNpe050-split2 > GCaMP8m. (D) DNpe050 (with Otd-FLP) activation elicits wing elevation in a small fraction of individuals. (i) A representative wing angle trace showing stable wing elevation during optogenetic activation. Dashed line: threshold wing angle criterion for wing elevation. (ii) Fraction of flies exhibiting sustained wing elevation (“penetrance”). (iii) Percentage of stimulation time performing wing elevation (“expressivity”). White and red shaded regions indicate quantifications before and during stimulation period, respectively. (E) Silencing DNpe050 (with Otd-FLP) using Kir2.1 reduces AIP-evoked wing elevation (i) and pumps (ii) but not turns (iii) or charges (iv). (F) Behavioral quantification for silencing of DNpe050 (with Otd-FLP) using Kir2.1 during natural threat display: (i) fraction of time performing wing elevation, (ii) average wing elevation bout duration, (iii) frequency of wing elevation, (iv) frequency of pumps, (v) frequency of turns during wing threat, (vi) frequency of charges during wing threat. Here and throughout, datapoints represent individual animals. Boxplots indicate median (box dividing line), interquartile range (box), and 1.5 times the interquartile range (whiskers). When less than 5 datapoints, lines denote median. Full genotypes of experimental flies for this and subsequent figures are listed in Table S1. Details of statistical analyses and sample size are given in Table S2. ns, not significant, *p < 0.05, **p < 0.01, ***p < 0.001, ****p < 0.0001, (*) pairwise significance that did not survive multiple comparison tests. For adjusted α values after multiple comparison corrections, see Table S2. See also Figure S2.

To confirm the connectome-predicted functional connectivity between AIP and DNpe050, we used two-photon microscopy to measure GCaMP fluorescence in DNpe050 during wide-field optogenetic activation of AIP neurons in head-fixed, living flies. Robust fluorescence increases were observed during photostimulation (Figure 2C), verifying that DNpe050 is functionally downstream and activated by AIP. The calcium response was absent in the genetic control group, which lacked the AIP driver but underwent the same LED illumination, suggesting that the observed response was not due to visual response to LED or leakage expression of Chrimson.

Next, we examined the behavioral function of DNpe050 by optogenetically activating the labeled cells in freely moving group-housed animals, which are non-aggressive and therefore do not exhibit spontaneous wing threat behavior. Under high-frequency (40 Hz) photostimulation, activation of DNpe050-split1 elicited stable wing elevation (defined by the criterion that both wings were elevated at between 35° and 65° for >0.2 s) (Figure 2Di), in approximately 10-15% of flies (Figure 2Dii). The penetrance (defined as the fraction of flies showing at least 1 instance of the behavior of interest during photostimulation) was consistent across various light intensities and frequencies with both DNpe050-split1 and DNpe050-split2 (Figure S2Ci, iii). Among responsive flies, the expressivity (defined as the percentage of the photostimulation period during which the behavior was performed) of wing elevation was highly variable but averaged approximately 10-40% (Figure 2Diii, and S2Cii, iv). Activation of DNpe050-split drivers did not elicit consistent changes in turning, charging, or pumping (Figure S2D).

To test the necessity of DNpe050 for AIP-driven behaviors, we performed epistasis experiments, in which AIP neurons were optogenetically activated with CsChrimson using a LexA driver, while DNpe050 was silenced using split-GAL4, Otd-FLP, and Kir2.1 effector. Silencing DNpe050 significantly suppressed AIP-elicited wing elevation (Figure 2Ei) and pumping (Figure 2Eii) but did not affect turning (Figure 2Eiii) or charging (Figure 2Eiv). Finally, we investigated whether DNpe050 is required for natural wing threat displays. Silencing DNpe050 using Kir2.1 and either of the two DNpe050-split drivers in pairs of single-housed male flies (which exhibit spontaneous wing threat behaviors) caused consistent reductions in the fraction of time flies performed wing elevation (Figure 2Fi), and in the average bout duration of wing elevation (Figure 2Fii). The frequency of wing elevation (Figure 2Fiii) and pump (Figure 2Fiv) were reduced in DNpe050-split1 but not or only mildly in split2. Turning (Figure 2Fv) and charging (Figure 2Fvi) during wing threat were not reduced. Lunging (Figure S2Ei) and general locomotion (Figure S2Eii) were not significantly different, suggesting that the reduction in wing elevation is not due to decreased overall aggressiveness or vigor.

Collectively, these results suggest that DNpe050 is sufficient to drive wing elevation - albeit with low penetrance - and is required for both wing elevation and pump during threat display.

### DNp60 elicits pump and wing elevation

We next investigated the function of DNp60, another DN that, like DNpe050, receives direct input from AIP (Figure 3A) and is predicted to be cholinergic by the connectome. Through screening combinations between hemi-drivers suggested by NeuronBridge (Figure S1C), we identified two split-GAL4 drivers for DNp60, DNp60-split1 (Figure 3B) and DNp60-split2 (Figure S3A). We again confirmed that DNp60 is functionally downstream of AIP by optogenetically activating the latter while imaging calcium transients in the former (Figure 3C).

**Figure 3.**
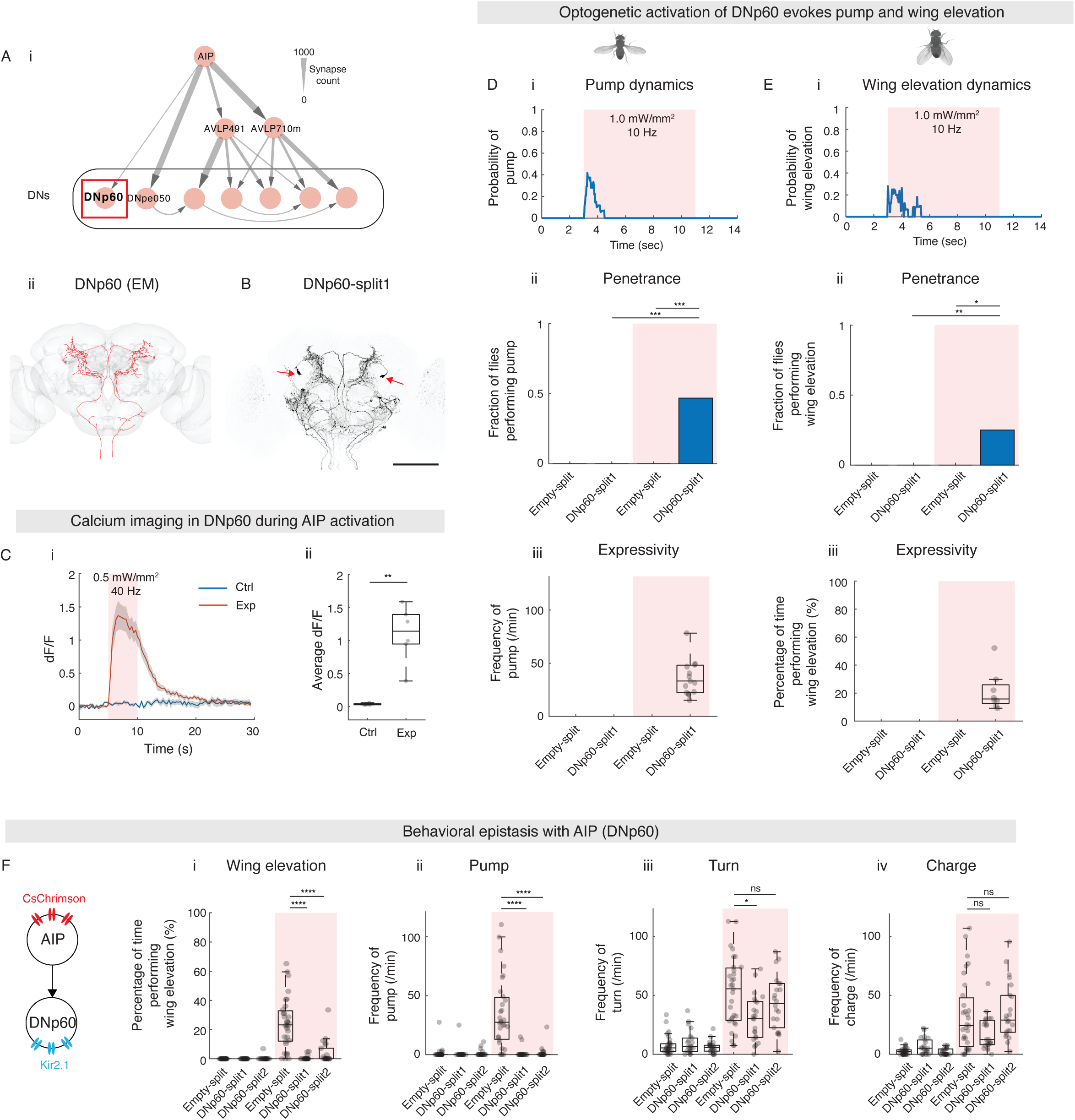
DNp60 elicits pump and wing elevation. (A) (i) DNp60 highlighted in the AIP downstream circuit diagram (red box). (ii) Cell skeletons of DNp60 reconstructed in the male CNS connectome. (B) Expression pattern of DNp60-split1. Arrows: cell body locations of DNpe050. Scale bar, 100μm. (C) jGCaMP8m fluorescence changes in DNp60 in response to AIP stimulation. (i) DNp60 response (dF/F) before, during, and after photostimulation of AIP. Red shaded region indicates light delivery. (ii) Average DNp60 response (dF/F) during photostimulation. Exp: AIP > Chrimson; DNp60-split1 > GCaMP8m. Ctrl: Empty > Chrimson; DNp60-split1 > GCaMP8m. (D) DNp60 (with Otd-FLP) activation elicits pump. (i) Probability of pump during a photostimulation trial. (ii) Fraction of flies performing at least one pump and (iii) frequency of pump during photostimulation. (E) DNp60 (with Otd-FLP) activation elicits wing elevation. Same as (D) but quantifications for wing elevation. (F) Silencing of DNp60 (with Otd-FLP) using Kir2.1 reduces AIP-evoked wing elevation (i) and pumps (ii) but not turns (iii) or charges (iv). See also Figure S3, Table S1 and S2.

Next, we investigated the behavioral phenotype elicited by DNp60 optogenetic activation. Medium-frequency (10 Hz) activation of DNp60 elicited pumps (Figure 3D), most of which occurred during the early phase of a trial (Figure 3Di). Pumps occurred in ∼50% of stimulated flies (Figure 3Dii), with a median frequency of ∼0.5 pumps/s per responsive fly. (Figure 3Diii). Higher LED frequencies and intensities did not further increase the penetrance or expressivity (pump count per responsive fly) (Figure S3Bi-ii). DNp60 activation also elicited wing elevation early in the stimulation period, with ∼25% penetrance and ∼20% expressivity (Figure 3E). Penetrance of wing elevation increased with LED frequencies and intensities, but saturated at ∼25%, while expressivity remained constant among responsive flies (Figure S3Biii-iv). Turning and charging were not elicited by DNp60 activation (Figure S3C).

To assess whether DNp60 is necessary for any AIP-induced wing threat actions, we conducted epistasis experiments in which DNp60 was silenced while AIP neurons were optogenetically activated. Silencing DNp60 significantly reduced AIP-elicited wing elevation (Figure 3Fi) and pump (Figure 3Fii) for both DNp60-split drivers. Turning was reduced only with DNp60-split1 but not with DNp60-split2 (Figure 3Fiii), likely due to off-target labeling. Charging was unaffected (Figure 3Fiv). Silencing DNp60 in pairs of interacting aggressive flies did not lead to any detectable reduction in natural wing threat actions (Figure S3D), perhaps reflecting redundancy with DNpe050.

Together, these results demonstrate that DNp60 is sufficient to elicit pump and wing elevation with moderate penetrance and low expressivity and is necessary for AIP-induced wing elevation and pump.

### DNpe050 and DNp60 synergistically control wing elevation and pump

Since DNpe050 activation drives wing elevation and DNp60 activation drives both pump and wing elevation, and each is necessary for AIP-induced wing actions, we hypothesized that coactivating DNpe050 and DNp60 in the same flies might enhance the penetrance and expressivity of wing elevation and pump.

To test this, we generated compound-genotype flies carrying both DNpe050-split1 and DNp60-split1 (Figure S4A) and compared the behavioral phenotypes of DNpe050 and DNp60 coactivation to those of single-neuron-type activation. At low frequency (2 Hz) photostimulation, DNpe050 activation alone triggered few stable wing elevation bouts (Figure 4Bi) and no pumps (Figure 4Ci). DNp60 activation alone elicited pumps and wing elevation, primarily during the initial light pulses of a trial (Figure 4Bii and 4Cii). In striking contrast, coactivation of DNpe050 and DNp60 induced rapid wing extension at each light pulse and maintained wings at an open angle between pulses (Figure 4Aiii), producing wing elevation that was sustained during the inter-pulse intervals (Figure 4Biii) and pumps across multiple photostimulation pulses throughout a trial (Figure 4Ciii). Coactivation significantly increased both penetrance (Figure 4Di) and expressivity (Figure 4Dii) of stable wing elevation relative to DNpe050 alone. Compared to DNp60 activation alone, coactivation showed equivalent pump and wing elevation penetrance (Figure 4Di and 4Ei), but significantly higher expressivity (Figure 4Dii and 4Eii). The expressivities of wing elevation and pump induced by coactivation exceeded the linear summation of those induced by single-neuron-type activation (Figure 4Dii and 4Eii, red dashed line). Importantly, coactivation of DNp60 and DNpe050 phenocopied AIP activation for both wing elevation and pump in terms of the magnitude of penetrance and expressivity (Figure 4D and 4E, AIP). The synergistic behavioral effect may result from reciprocal connectivity between DNpe050 and DNp60, or from their convergent input onto common downstream targets (see Discussion).

**Figure 4.**
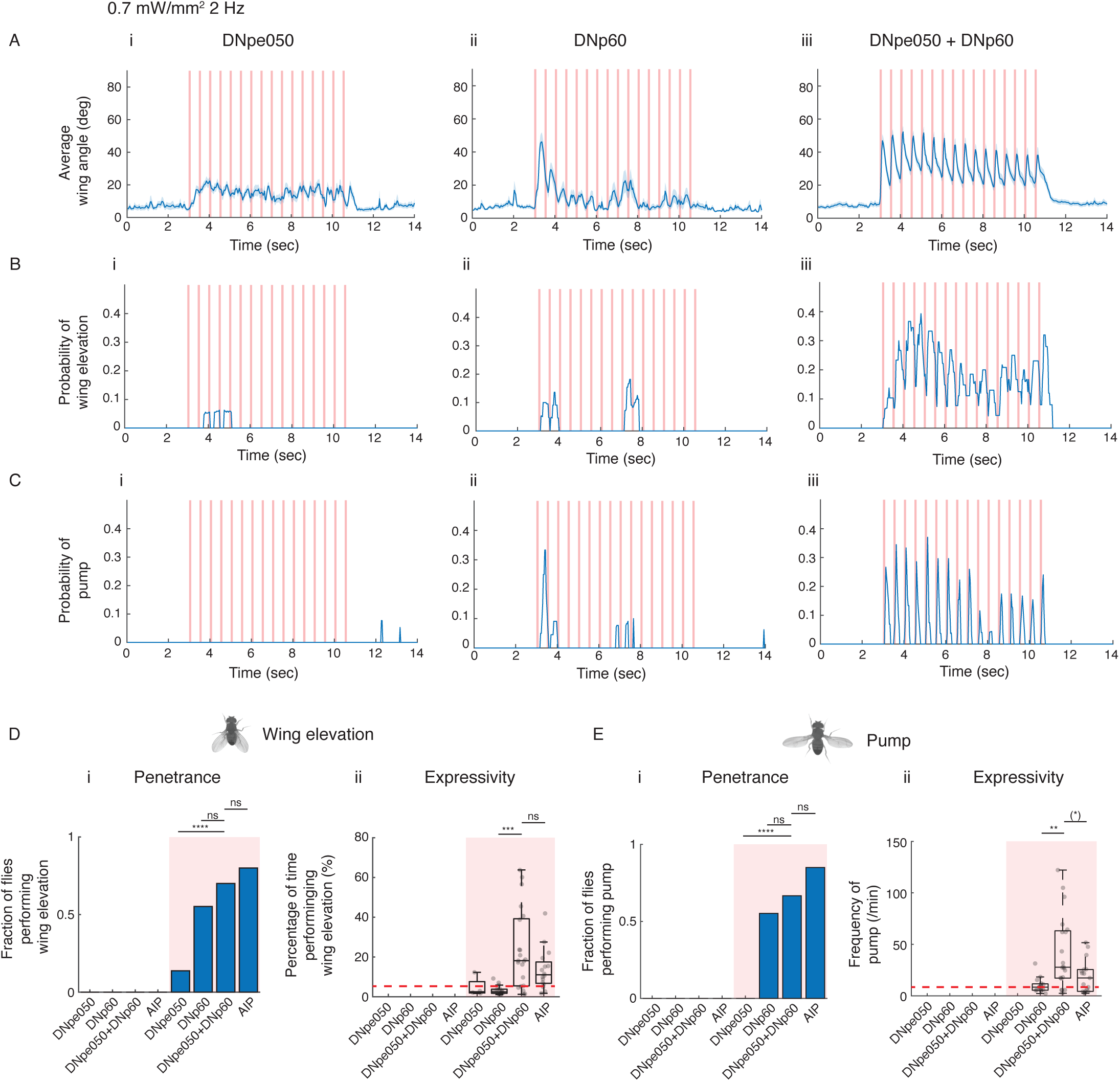
DNpe050 and DNp60 synergistically control wing elevation and pump. (A) Wing angles before, during, and after photostimulation of (i) DNpe050-split2 (with Otd-FLP), (ii) DNp60-split1 (with Otd-FLP), and (iii) DNpe050-split1 and DNp60-split2 combined (with Otd-FLP, effectively labeling DNpe050-split2 neurons as well, see Table S1 for detailed genotypes). Here and throughout, shaded lines denote mean ± SEM. (B) Probability of wing elevation during photostimulation of (i) DNpe050 (with Otd-FLP), (ii) DNp60 (with Otd-FLP), and (iii) DNpe050 and DNp60 combined (with Otd-FLP). (C) Probability of pump during photostimulation of (i) DNpe050 (with Otd-FLP), (ii) DNp60 (with Otd-FLP), and (iii) DNpe050 and DNp60 combined (with Otd-FLP). (D) Fraction of flies performing at least one wing elevation bout (i) and percentage of time performing wing elevation (ii) during photostimulation. Red dashed line indicates the linear summation of medians of DNpe050 and DNp60 activation alone. (E) Fraction of flies performing at least one pump (i) and frequency of pump (ii) during photostimulation. Red dashed line indicates the linear summation of medians of DNpe050 and DNp60 activation alone. See also Figure S4, Table S1 and S2.

Under high frequency (40 Hz) photostimulation, coactivation similarly increased penetrance (Figure S4Bi) and expressivity (Figure S4Bii) of wing elevation relative to either driver alone. Pump was less effectively elicited by activation of any driver tested at 40 Hz, including that of AIP. This ceiling effect likely precluded a significant coactivation improvement for pump (Figure S4Biii and S4Biv). To rule out the possibility that the observed behavioral effects were due to off-target neuron activation or synergistic effects between off-targets neurons and DNpe050/DNp60, we repeated the coactivation experiments using DNp60-split2 combined with four different DNpe050 split-GAL4 lines, each with distinct off-target patterns. Similar results were obtained under both 2 Hz (Figure S4C) and 40 Hz stimulation (Figure S4D).

Taken together, these data suggest that DNpe050 and DNp60 act synergistically to promote wing elevation and pump, and their coactivation can recapitulate the wing actions driven by AIP.

### AVLP491 induces locomotion and is necessary for AIP-elicited threat locomotor actions

We focused next on the function of INs AVLP710m and AVLP491 (Figure 5A and 6A), both of which receive direct synaptic input from AIP neurons and in turn project to multiple DNs. Based on the connectome, AVLP491 is predicted to be cholinergic. To investigate the function of AVLP491, we identified two split-GAL4 drivers, AVLP491-split1 (Figure 5B) and AVLP491-split2 (Figure S5A). Calcium imaging confirmed that AVLP491 is functionally downstream of AIP (Figure 5C), consistent with the projection suggested by the connectome.

**Figure 5.**
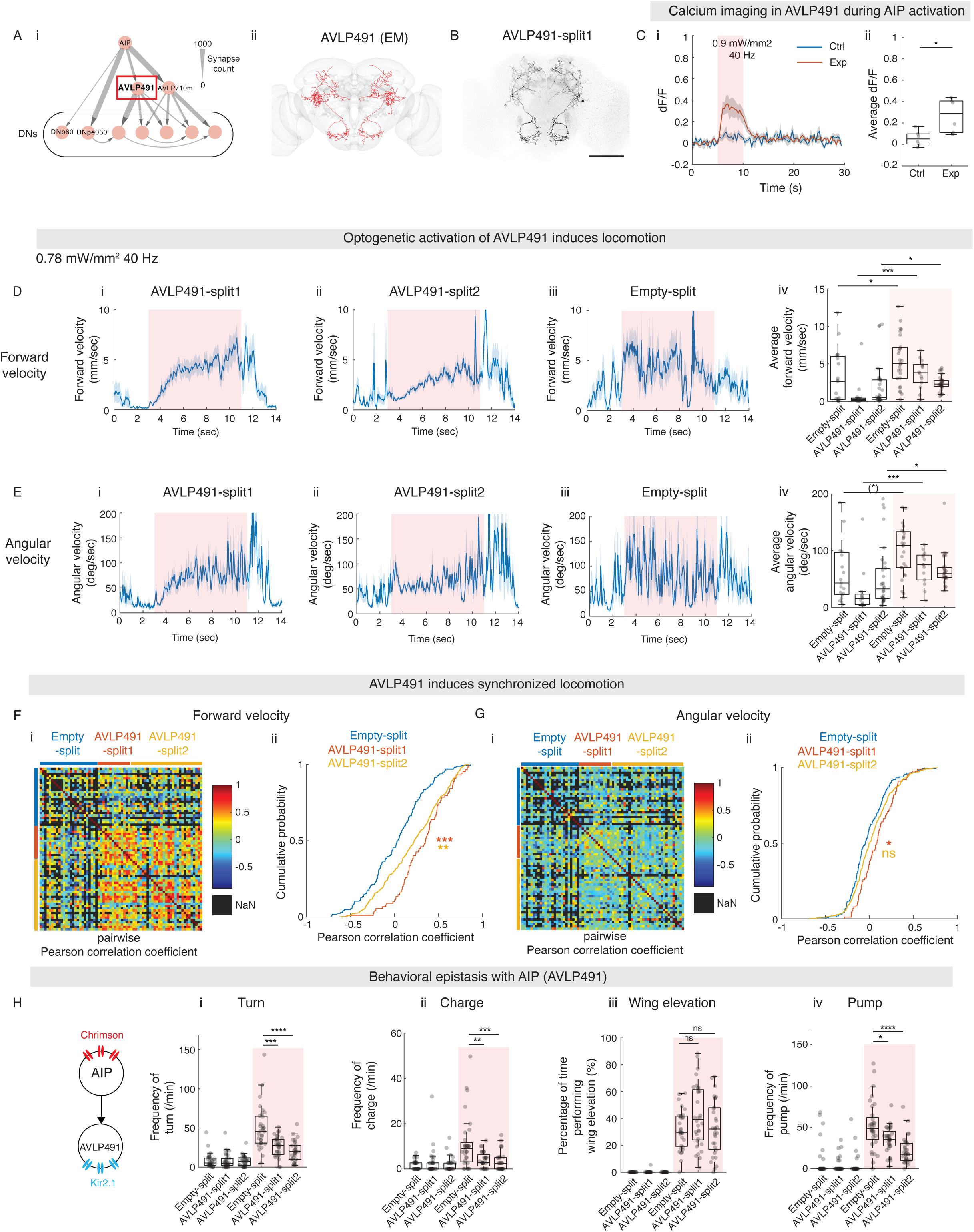
AVLP491 induces locomotion and is necessary for AIP-elicited threat locomotor actions. (A) (i) AVLP491 highlighted in the AIP downstream circuit diagram (red box). (ii) Cell skeletons of AVLP491 reconstructed in the male CNS connectome. (B) Expression pattern of AVLP491-split1 (with Otd-FLP). Scale bar, 100μm. (C) jGCaMP8m fluorescence changes in AVLP491 in response to AIP stimulation. (i) AVLP491 response (dF/F) before, during, and after photostimulation of AIP. Red shaded region indicates light delivery. (ii) Average AVLP491 response (dF/F) during photostimulation. Exp: AIP > Chrimson; AVLP491-split1 > GCaMP8m. Ctrl: Empty > Chrimson; AVLP491-split1 > GCaMP8m. (D) Forward velocities before, during, and after optogenetic activation of AVLP491-split1 (with Otd-FLP) (i), AVLP491-spilt2 (with Otd-FLP) (ii), and Empty-split (with Otd-FLP) (iii). (iv) Average forward velocities before (white) and during photostimulation (red shaded). (E) Same as (D) but for angular velocities. (F) (i) A heatmap showing pairwise Pearson correlation coefficients of forward velocities during photostimulation period. Each row and column indicate an individual fly. NaN: not a number, when data not qualified (see Methods) (ii) Cumulative distribution of pairwise Pearson correlation coefficients displayed in (i). (G) same as F but for angular velocities. (H) Silencing of AVLP491 reduces AIP-elicited turns (i), charges (ii), and pumps (iv) but not wing elevation (iii). See also Figure S5, Table S1 and S2.

**Figure 6.**
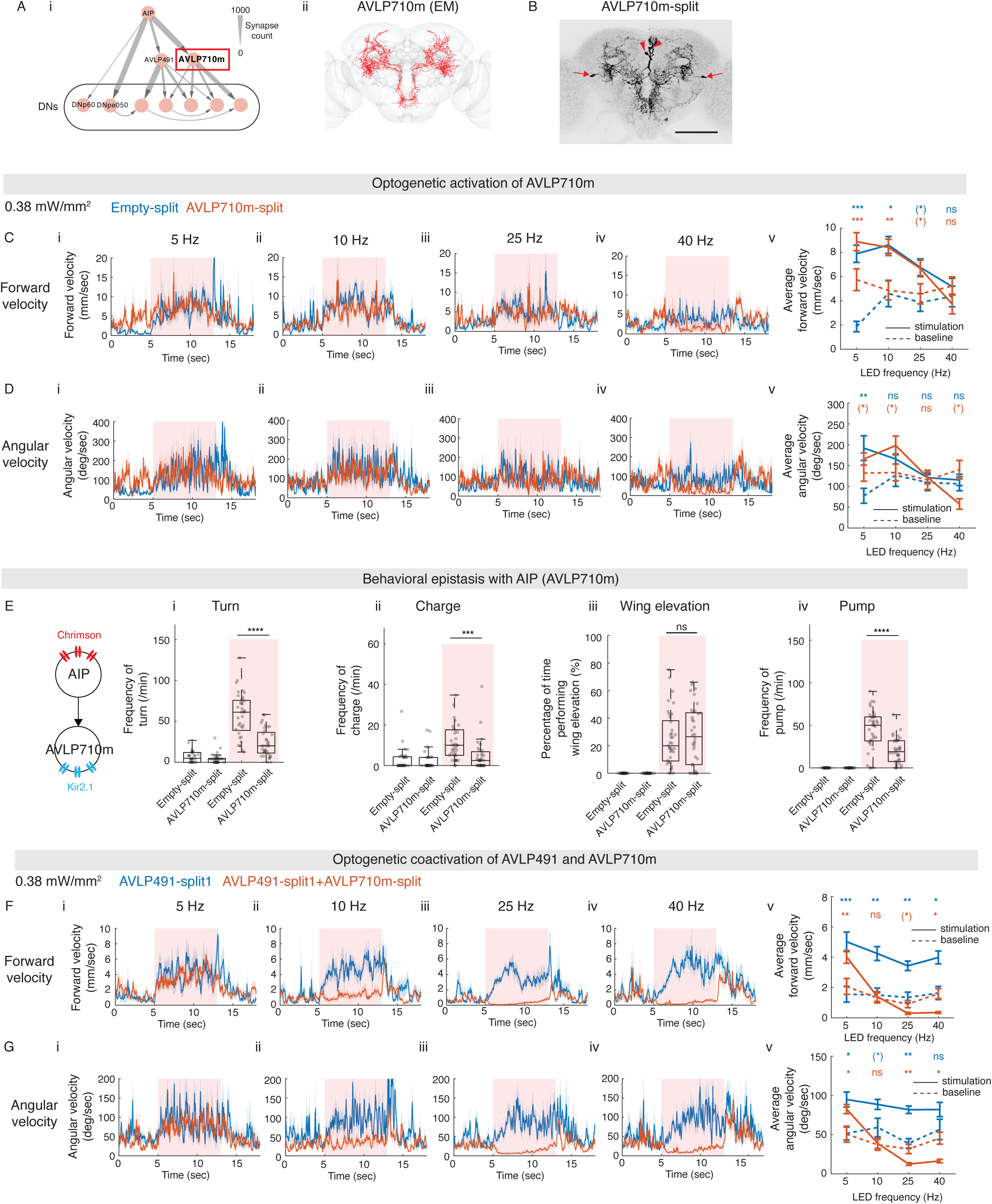
AVLP710m functions antagonistically to AVLP491 to control locomotion. (A) (i) AVLP710m highlighted in the AIP downstream circuit diagram (red box). (ii) Cell skeletons of AVLP710m reconstructed in the male CNS connectome. (B) Expression pattern of AVLP710m-split. Arrows: cell body locations of AVLP710m. Arrowheads: cell body locations of off-target neurons. Scale bar, 100μm. (C) Forward velocities before, during, and after optogenetic activation of AVLP710m (with Otd-FLP) at 5 Hz (i), 10 Hz (ii), 25 Hz (iii) and 40 Hz (iv). (v) Average forward velocities before (dashed line) and during (solid line) photostimulation. Statistic comparisons are performed within genotypes between before (dashed line) and during (solid line) photostimulation. Lines and error bars denote mean ± SEM. (D) same as (C) but for angular velocities. (E) Silencing of AVLP710m reduces AIP-elicited turns (i), charges (ii), and pumps (iv) but not wing elevation (iii). (F) Forward velocities before, during, and after optogenetic activation of AVLP491 (with Otd-FLP) alone or AVLP491 and AVLP710m combined (with Otd-FLP) at 5 Hz (i), 10 Hz (ii), 25 Hz (iii) and 40 Hz (iv). (v) Average forward velocities before (dashed line) and during (solid line) photostimulation. Statistic comparisons are performed within genotypes between before (dashed line) and during (solid line) photostimulation. (G) same as (F) but for angular velocities. See also Figure S6, Table S1 and S2.

Optogenetic activation of AVLP491 induced locomotion and in freely moving flies, with increased forward velocity (Figure 5Di-ii, iv) and angular (Figure 5Ei-ii, iv) velocity, respectively, relative to pre-stimulation baseline. Under some conditions a gradual increase in velocity was observed throughout the photostimulation period. Locomotion induced by AVLP491 activation was also temporally correlated across individuals. To quantify this, we calculated pairwise Pearson correlation coefficients for forward or angular velocities across individuals of all genotypes (Figure 5F and 5G). For forward velocity, AVLP491-activated flies showed high correlation among flies with the same split-Gal4 driver, as well as across both AVLP491-split drivers (Figure 5Fi), and this correlation was significantly higher than that observed in the genetic control (Figure 5Fii). For angular velocity, both AVLP491-split drivers displayed a higher correlation than did controls but the effect was statistically significant only for AVLP491-split1 (Figure 5Gi, ii), potentially reflecting the more variable nature of orienting behavior. Although genetic controls lacking opsin expression also showed increased forward velocity (Figure 5Diii, iv) and angular velocity (Figure 5Eiii, iv), the locomotor response in response to photostimulation in control appeared irregular and lacked the temporal correlation (Figure 5F and 5G) observed in the AVLP491 genotypes, perhaps reflecting the fact that sudden bursts of illumination can induce a locomotor startle response. Notably, activating AVLP491 did not induce any wing elevation or pump (Figure S5B). Therefore, AVLP491 activation specifically induced locomotion.

In epistasis experiments, silencing AVLP491 significantly suppressed AIP-induced turning and charging (Figure 5Hi and 5Hii), indicating a requirement of these neurons for leg-mediated threat actions. Epistatic silencing of these cells did not reduce AIP-evoked wing elevation (Figure 5Hiii). However, we observed a significant reduction in pump (Figure 5Hiv). This effect could be a direct result of AVLP491 silencing, or could also be an indirect effect due to reduced turning and charging, because pump is frequently accompanied by turns and charges^16^, suggesting that the execution of pump might require proprioceptive feedback from the legs. Silencing AVLP491 in pairs of spontaneously aggressive flies using either of the two split-GAL4 drivers did not lead to any consistent reduction in threat actions, including turning or charging (Figure S5C), perhaps reflecting redundancy among locomotion-related circuits activated in this naturalistic setting.

Together, these data suggest that AVLP491 is sufficient to drive locomotion and is necessary for AIP-elicited turning and charging.

### AVLP710m functions antagonistically to AVLP491

AVLP710m is the only other major IN target of AIP besides AVLP491 that projects directly to DNs (Figure 6A). We identified a split-GAL4 driver (Figure 6B) for AVLP710m. Fluorescence *in situ* hybridization with HCR3.0^39^ revealed that AVLP710m expresses VGAT but not ChAT (Figure S6A), indicating it is an inhibitory interneuron. Although the connectome suggests over 1,000 synapses between AIP and AVLP710m (Figure 6A), we did not detect significant calcium responses in AVLP710m when AIP was photostimulated in head-fixed preparations (data not shown). This may be because AVLP710m is sensitive to the state of the fly and could not be successfully activated by AIP under restrained open-cuticle conditions. Consistent with this, AVLP710m receives in total around 15,000 input synapses from over 500 other neurons (with a cutoff at 5 synapses), which may include tonic inhibitory input that overrides AIP activation.

Next, we used direct optogenetic activation of AVLP710m to probe its behavioral function. At an LED intensity of 0.38 mW/mm^2^ and frequencies of 5, 10, and 25 Hz, the experimental flies did not display any differences from the genetic controls (Figure 6Ci-iii, v and 6Di-iii, v). At the highest frequency tested (40 Hz), AVLP710m activation caused a trend towards reduced forward and angular velocity during photostimulation, relative to pre-stimulation baseline, but this effect failed to reach statistical significance (Figure 6Cv and 6Dv). However, when the LED intensity was increased to 0.78 mW/mm^2^ at 40 Hz, stimulation of AVLP710m significantly reduced both forward and angular velocity (Figure S6B), suggesting that AVLP710m can suppress light-evoked general locomotion only when strongly activated.

To test AVLP710m’s necessity for AIP-evoked threat behaviors, we silenced it in epistasis experiments. AVLP710m silencing using Kir2.1 significantly suppressed AIP-induced turning (Figure 6Ei), charging (Figure 6Eii), and pump (Figure 6Eiv), but did not affect wing elevation (Figure 6Eiii). This suggests that although direct AVLP710m activation can suppress general locomotion, it is necessary for AIP-evoked transient, discrete bouts of turning and charging. Silencing AVLP710m in single-housed males performing spontaneous threat behavior did not reduce natural turning and charging (Figure S6Cv, vi), probably reflecting redundancy among locomotion-related circuits or insufficient inhibition. A mild reduction in wing elevation was observed (Figure S6Ci, ii).

Given that both AVLP491 and AVLP710m influence locomotion and are required for AIP-driven turning and charging, we hypothesized that they may work together to generate threat locomotor actions. To test this hypothesis, we performed coactivation experiments. AVLP491 activation alone evoked increased locomotion at all frequencies tested (Figure 6F and 6G). However, at 10 Hz or above, coactivation with AVLP710m abolished this increase (Figure 6Fii-v and 6Gii-v). Furthermore, at 25 Hz or 40 Hz, in coactivated flies forward and angular velocities dropped below pre-stimulation baseline (Figure 6Fiii-v and 6Giii-v), approaching zero. Similar results were obtained with AVLP491-split2 (Figure S6D and S6E). Given that AVLP710m has a high stimulation threshold for suppressing general locomotion (Figure 6C, 6D, and S6B, 0.78 mW/mm^2^, 40 Hz), but a much lower threshold for suppressing AVLP491-induced locomotion (Figure 6F, 6G, 0.38 mW/mm^2^, 10 Hz), these results suggest a specific antagonistic interaction between AVLP491 and AVLP710m. Consistent with this notion, connectomic data show that ALVP491 and AVLP710m project to overlapping DNs (Figure S6F), including DNg100 and DNge050, which are functionally implicated in locomotion^40,41^. This suggests that their functional antagonism may result from converging input onto common direct downstream targets.

Taken together, these data indicate that AVLP710m is necessary for AIP-induced turning and charging and functions antagonistically to AVLP491 to regulate locomotion.

### Temporally patterned AVLP491 and AVLP710m coactivation recapitulates AIP-induced turn and charge-like behavior

Turns and charges in the context of threat are performed as transient, discrete bouts of locomotion^16^. AIP activation mimics this pattern, eliciting brief, punctuated turns and charges, even during constant LED stimulation (Figure 7A). In contrast, AVLP491 activation alone leads to sustained locomotion (Figure 5D and 5E), while its coactivation with AVLP710m leads to continuous immobility (Figure 6F, 6G, S6D and S6E). Despite these contrasting results, both AVLP491 and AVLP710m are required for AIP-induced turning and charging (Figure 5H and 5F). How, then, does AIP generate transient, discrete locomotor bouts through these neurons?

**Figure 7.**
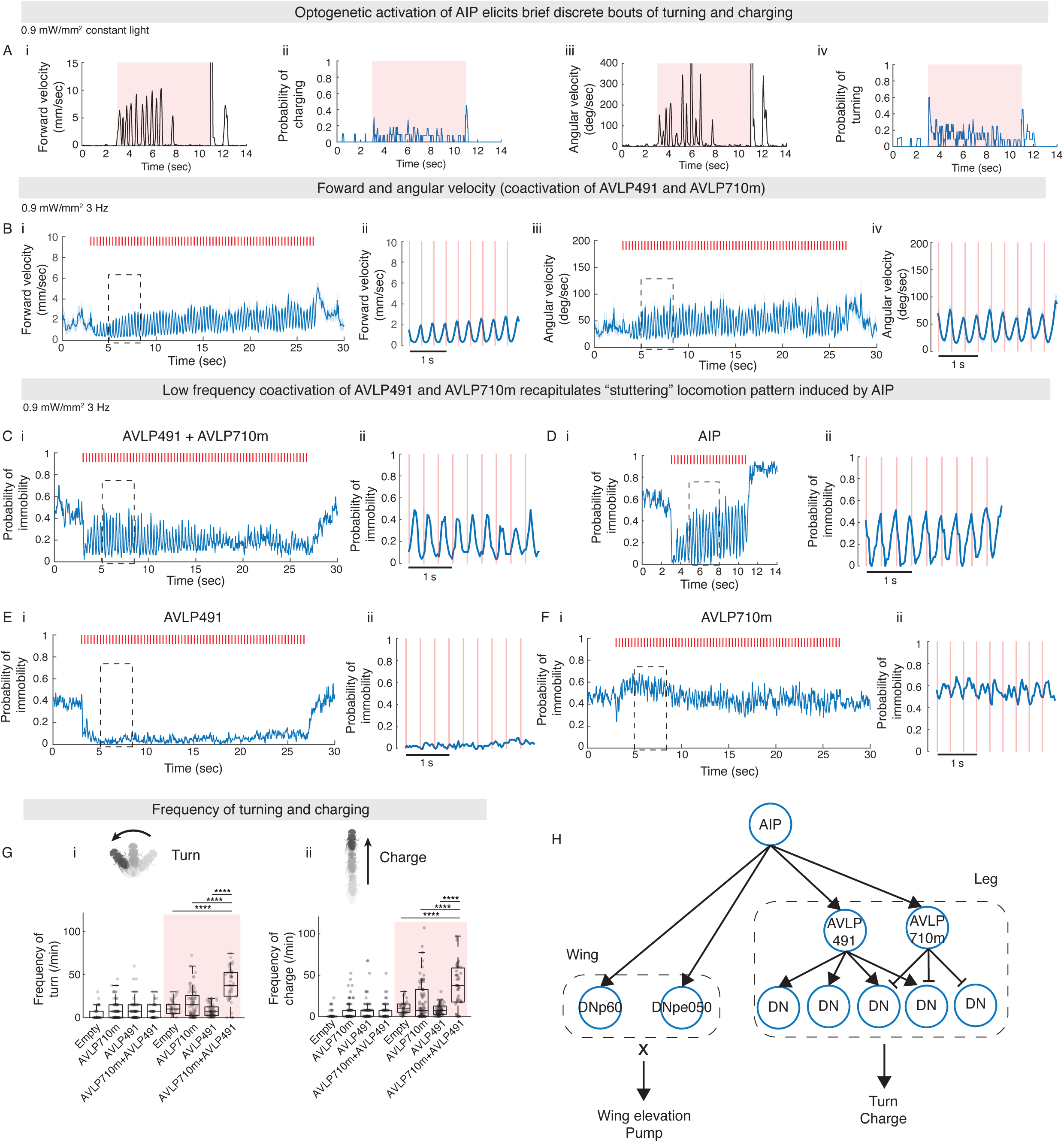
Temporally patterned AVLP491 and AVLP710m coactivation recapitulates AIP-induced turn and charge-like behavior. (A) Forward (i) and angular (iii) velocity traces of an example fly during AIP (with Otd-FLP) photostimulation with constant LED light. Probability of charging (ii) and turning (iv) during AIP (with Otd-FLP) photostimulation with constant LED light. (B) Forward (i) and angular (iii) velocity before, during, and after photostimulation. (ii) and (iv), insets (dashed box) of (i) and (iii) respectively from second 5 to 8. Red bar indicates light pulse delivery. (C) (i) Probability of immobility before, during, and after coactivation of AVLP491 and AVLP710m (with Otd-FLP). (ii), inset of (i) from second 5 to 8. (D-F) same as (C) but for activation of AIP (with Otd-FLP) (D), AVLP491 (with Otd-FLP) (E), and AVLP710m (with Otd-FLP) (F). (G) Quantification of turning (i) and charging-like behavior (ii) during photostimulation (with Otd-FLP). (H) A working model of AIP downstream circuit See also Table S1 and S2.

We hypothesized that while AIP activities provide overall initiation and coordination of threat behavior, AVLP491 and AVLP710m may shape the temporal structure of locomotor bouts. If true, their activation frequency should match the dynamics of turning and charging, which is constrained by the physical limitation of motor execution. We therefore tested this by coactivating AVLP491 and AVLP710m at 3 Hz, a frequency comparable to the turning and charging frequency observed during AIP activation under constant light (Figure 7A). Surprisingly, coactivation at 3 Hz produced oscillatory patterns in both forward (Figure 7Bi-ii) and angular velocities (Figure 7Biii-iv).

The turns and charges observed during both natural and AIP-evoked threat are exhibited as brief bouts of forward and angular locomotion, interspersed with periods of immobility^16^. Therefore, we quantified the probability of immobility (defined as periods during which forward velocity is < 1mm/s and angular velocity is < 10°/s) during coactivation of AVLP491 and AVLP710m. Strikingly, in flies coactivated at 3 Hz, bouts of forward and angular locomotion were interspersed with regular periods of immobility (Figure 7C). This “stuttering” locomotion pattern closely resembled the turns and charges elicited by AIP activation at the same frequency (Figure 7D) and was notably absent when either IN was activated alone (Figure 7E and 7F). Quantification of turn- and charge-like bouts showed that coactivation of AVLP491 and AVLP710m significantly increased the frequency of these actions relative to AVLP491 or AVLP710m alone (Figure 7G).

In summary, low-frequency coactivation of AVLP491 and AVLP710m recapitulated the temporal pattern of AIP-induced threat locomotion, generating discrete bouts of locomotion not observed with activation of either IN alone. These findings suggest that AVLP491 and AVLP710m act in combination to generate the threat locomotor actions mediated by the legs.

## Discussion

How neural circuits organize and coordinate the movements of different appendages during social displays is poorly understood. In this study, we identified a hierarchical circuit containing four newly characterized cell types that collectively mediate the actions comprising threat displays evoked by stimulation of AIP neurons (Figure 7H). DNpe050 and DNp60, two direct DN targets of AIP, synergistically control wing elevation and pump, while AVLP491 and AVLP710m, two functionally antagonistic INs projecting to overlapping but distinct populations of DNs, coordinatively control threat locomotor actions, i.e., rapid orientation (turning) and acceleration (charging) towards the opponent fly. These findings suggest that appendicular control during threat display, as driven by AIP activation, is mediated by specialized downstream functional neural modules that use combinatorial coding to coordinate distinct wing- and leg-mediated actions. They also reveal a fundamentally different circuit logic from the simpler models that intuition might suggest (Figure 1B).

### A hierarchical circuit for threat action control

Nicholas Tinbergen proposed a conceptual framework in which the neural control of innate behaviors is organized hierarchically^42–44^, with reproductive instincts controlled at the highest level, activities such as aggression, nesting and mating at the second level, and specific aggressive behaviors such as biting, chasing, and threatening at the third level (Figure S7A). AIP appears to correspond to a “third-level center” in this framework.^16^ Third-level centers were proposed in turn to control specific appendages, followed by appendicular substructures, their muscles and motor units. However, very few neural systems have been studied at these more granular levels of behavioral control. Therefore, it is not clear whether these lower levels are in turn hierarchically organized or not. Our findings show that AIP neurons hierarchically exert control over a group of DNs that convey information from the central brain to the VNC (Figure 7H). However, rather than conforming to a simple one-layer fan-out design (Figure 1Bii), AIP downstream circuits comprise both direct projections to DNs and an indirect IN->DN sub-hierarchy. Moreover, the neural control of behavior by these lower-level centers is combinatorial. Thus, the hierarchical control of behavior is deeper and more granular, as well as more complex in its neural implementation, than Tinbergen envisioned.

### Appendage-specific modules for action control

AIP output is organized hierarchically into two different appendage-specific modules: one controlling wing actions and the other regulating leg actions. The former actions are mediated by direct projections to DNs, while the latter are mediated by indirect projections via an intervening layer of INs. This difference in circuit organizational logic likely reflects different neural control requirements for these two types of appendages.

Wing threat actions are relatively stereotyped. During wing elevation, wings typically are raised to a fixed 45° angle and remain there. During a pump both wings are briefly expanded to approximately 90°. Thus, unlike flying or singing which involves flexible and dynamic control of the wings^45,46^, threat displays involve “one-shot” changes in wing postures. Such stereotyped wing postures may require fast, reliable triggering by dedicated DNs, making direct monosynaptic AIP-to-DN connectivity advantageous.

In contrast to wing movements during threat, locomotion is prevalent in various behavioral contexts and can exhibit different modes^47–52^. Flies can walk backwards^48,49^, follow a target^47,51^, walk straight^47^, pivot and swerve^50^, which can be driven by single neuronal types or by the coordinated activity of multiple neuronal types. DNs controlling the legs may participate in a broad set of walking-related motor programs. The presence of an intermediate IN layer downstream of AIP allows for two key forms of circuit control: 1) recruiting the appropriate DN population to produce a motor action specific for a particular behavioral display – a “corralling” function – and 2) shaping the temporal dynamics of DN activity to generate appropriately timed and structured outputs – a “sculpting” function. Although the four types of threat actions are relatively stereotyped, turning and charging are inherently dynamic, with direction, speed, and bout duration modulated by the fly’s orientation and distance relative to its opponent. Their execution likely depends on real-time sensory input to guide and adjust the trajectory of the movement. The layered AIP-IN-DN architecture could facilitate these dynamics by allowing sensory signals to modulate circuit activity at multiple stages.

Overall, segregating wing and leg control into sub-modules specialized for threat enables tailored neural strategies, ensuring precise and efficient execution of actions with different functional demands while still coordinating these actions into an ethologically meaningful communicative display.

### Combinatorial coding of actions

A main and surprising finding of our analysis is that individual downstream target neurons of AIP are not dedicated to single threat actions, as one might have assumed (Figure 1B). Instead, they function in a combinatorial manner to control appendage-specific actions.

In the wing module, DNpe050 and DNp60 act synergistically to promote both wing elevation and pumping. When activated individually, each neuron elicits one or both actions, but with low penetrance and expressivity. In contrast, their coactivation significantly enhances expressivity of pumping and the penetrance and expressivity of wing elevation. In the central brain, connectomic data suggest that there is no direct connection between DNpe050 and DNp60. In the VNC, we identified only one high-ranking direct downstream target (IN19B007) shared by DNpe050 and DNp60 (Figure S7Bi). However, there is extensive interconnectivity between the first-order downstream targets of these two DNs in the VNC (Figure S7Bii), suggesting that the synergy might happen at this level. Regardless of the details of implementation, this functional synergy suggests an arithmetic “multiplication” computation, whereby the combined activation of the two DNs produces a non-linear behavioral output that is stronger and/or more reliable than a linear combination of their individual behavioral effects.

In contrast to the direct activation of DNs for wing threat movements, DNs controlling turning and charging (Figure S6F) are regulated by two antagonistic INs with opposite influences: AVLP491 is excitatory and promotes locomotion, while AVLP710m is inhibitory and can effectively and specifically suppress AVLP491-elicited locomotion. This antagonism aligns with arithmetic “subtraction” in which excitatory drive is counterbalanced by targeted inhibition to regulate locomotor output. Thus, combinatorial control of the two appendage-specific modules downstream of AIP neurons involves apparent multiplication and subtraction-like behavioral computations, respectively. Where and how these computations are implemented will be an interesting subject for further studies.

How DNs are organized to convey motor instructions from the brain to the VNC remains a topic of debate^53,54^. While some DNs function as command-like neurons^47,48,55–65^, others may operate through population coding^66–69^. Recent studies suggest organizational principles by which actions are driven by networks of DNs organized under superordinate command-like DNs^53^ or through nested DN activities^70,71^. Our findings introduce two novel strategies by which DNs may control motor actions – “multiplication” between two DNs and “subtraction” by a pair of INs that directly control multiple DNs. DNs have been considered as an information bottleneck between the brain and the VNC because they are greatly outnumbered by other types of neurons in both locations. This combinatorial coding strategy thus could enable greater complexity and adaptive action control given a limited number of DNs.

### Generation of turn- and charge-like locomotion patterns

AIP elicits transient, discrete bouts of turning and charging, even under constant LED stimulation. In contrast, high-frequency coactivation of AVLP491 and AVLP710m results in immobility; only low-frequency coactivation produces brief turn- and charge-like locomotor bouts (“stuttering”) that resemble the behavioral effect of AIP activation. This raises a key new question: how is continuous AIP activation transformed into discrete, brief turning or charging bouts through AVLP491 and AVLP710m? One possibility is that the synaptic properties between AIP and these two IN targets may define a transfer function that intrinsically limits their firing patterns. Specifically, AIP may activate AVLP710m with a temporal delay relative to AVLP491, so that AVLP491-elicited extended locomotion is truncated into brief bouts of discrete locomotion. In contrast to the effect of AIP activation, we observed a phase shift in the timing of locomotion relative to LED pulses in AVLP491/AVLP710m coactivation flies, suggesting that rebound excitation in downstream locomotor control circuits may also be involved. Alternatively, other cellular or circuit mechanisms may generate intermittent pulsatile output from AIP neurons despite continuous photostimulation. While future investigation is required to test these possibilities, our findings suggest that precise timing of AVLP491 and AVLP710m activities are critical for turning and charging.

### Flexible coupling and decoupling of actions under a hierarchical circuit

Threat actions can be flexibly performed both independently and coordinately. While wing elevation can occur with or without turns and charges, pumping is typically accompanied by these locomotor actions^16^. Our data reveal how a threat display which coordinates both sets of appendages is achieved via AIP recruitment of both wing and leg modules, in a hierarchical manner. However, this organization raises a key question: how does this circuit enable both coupling and decoupling of threat actions? While in principle this might have been achieved using separate subpopulations of AIP neurons to control the wings and legs, our connectomic analysis reveals that this is not the case.

One possibility is that different levels of AIP activity may govern action selection. Our initial characterization of AIP neurons indicated that optogenetic activation of these neurons can elicit threat actions in a scalable manner, with turning and charging requiring a lower threshold than wing elevation and pump^16^. The stereotyped co-occurrence of pumping with turn and charge could be explained by this threshold model: when AIP activity reaches the level required to trigger pumping, it has already reached the activation thresholds for turning and charging, ensuring their simultaneous execution.

However, this mechanism alone cannot explain how wing elevation can occur without turning and charging^16^, since the latter have a lower activation threshold than the former. In the absence of additional control mechanisms, a simple ramp-to-threshold model would predict that all instances of wing elevation should be accompanied by turning and charging, and this is not observed. Several potential mechanisms could explain this uncoupling. First, once AIP activity reaches the threshold for eliciting wing elevation, higher levels may recruit an inhibitory circuit node that evokes immobility. This node could be the GABAergic AVLP710m; DNpe050/DNp60, via their downstream circuits in the VNC; or another, as-yet uncharacterized downstream target. Second, proprioceptive feedback from wing elevation may actively suppress locomotor actions under certain conditions. Third, locomotion during social behaviors may be modulated by external sensory cues^72–75,40^. Turn or charge could be suppressed when the fly is already facing its opponent or in close proximity, preventing unnecessary movements. Further studies will be required to distinguish these alternatives.

### Potential generalizability of AIP downstream circuit logic

The hierarchical, body-part-specific, combinatorial organization we describe for threat displays may generalize to other innate behaviors requiring coordinated use of multiple body parts – such as walking, grooming^56,57,76^, hunting, or escape^77,78^. For instance, coordinated leg-wing actions during escape involve giant fiber DNs that directly excite leg extensor motor neurons and wing depressor interneurons^78^. However, our data reveal a more complex and flexible circuit that allows separate while cohesive activation of movements performed by different body parts during a social display. This is more analogous to what is observed during courtship, where singing can be performed both by stationary male flies as well as during chasing or circling a female^45^. Similar circuit motifs might underlie coordinated multi-appendicular behaviors across species.

### Limitations of this study

Despite the insights provided by this study, several limitations should be acknowledged. First, although we were able to functionally elicit or inhibit threat actions through neuronal activity perturbation, we could not directly record the activity of AIP neurons or their downstream targets during natural wing threat displays. Achieving this would require an imaging setup in which wing threat actions can be evoked from flies by naturalistic visual stimuli in a head-fixed open-cuticle preparation under 2-photon optics – a technically challenging preparation that we were unable to establish despite extensive efforts. Second, in this study, we analyzed the function of four different neuron types that are direct synaptic targets of AIP neurons, but the roles of other targets, or indirect DN targets of AIP remain uncharacterized. A comprehensive understanding of the subtleties of wing threat control will necessitate analysis of these additional cell types, alone and in combination with those characterized here. Such analyses may provide further insight into the interactions among AIP downstream targets in behavioral control, potentially explaining the lack of loss-of-function phenotypes observed in DNp60, AVLP491, and AVLP710m silencing during natural wing threat, as well as suggest how sensory cues may dynamically regulate action selection.

## Supplemental Figure Legends

**Figure S1.**
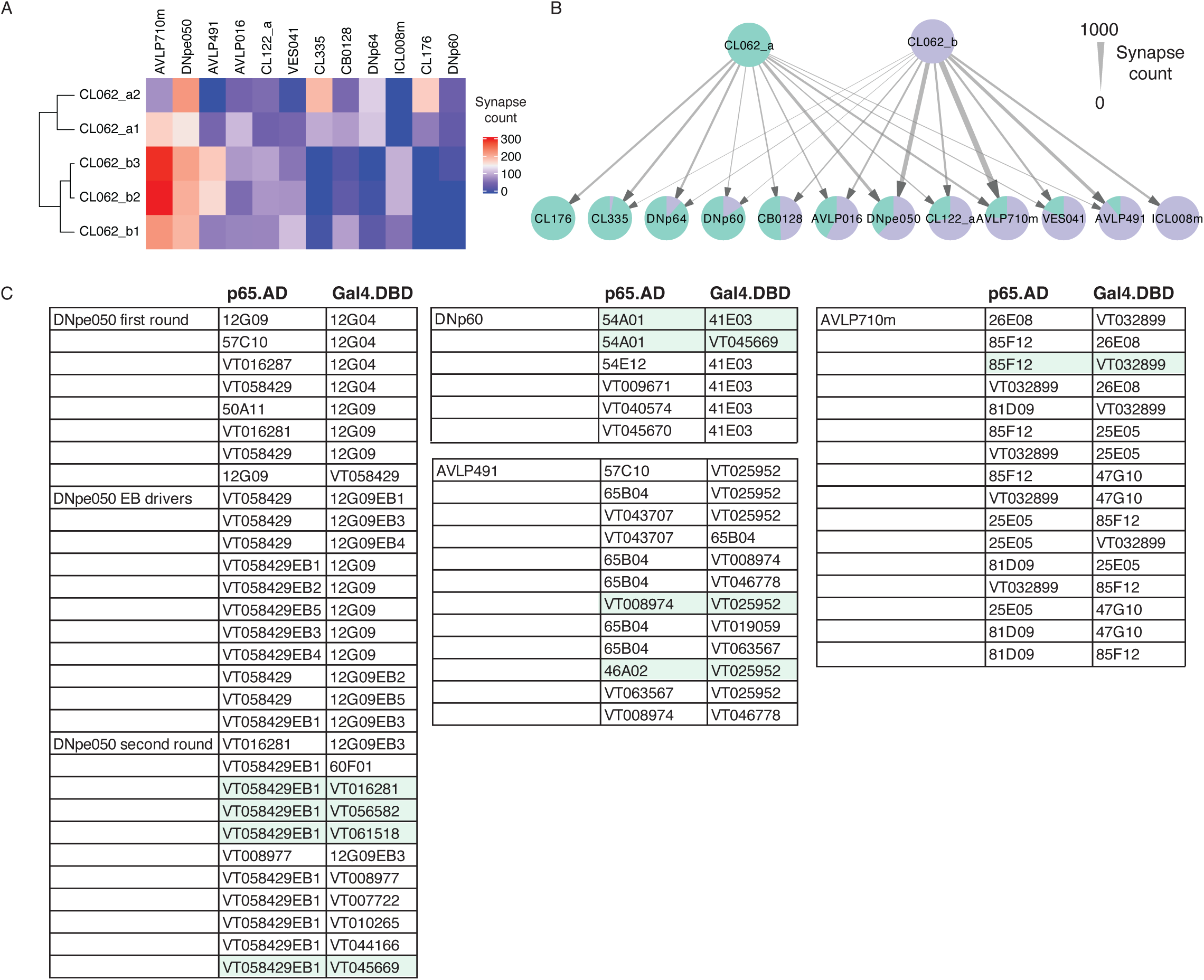
Heterogeneity in AIP population and other AIP downstream targets. (A) A heat map of synapse counts between individual pairs of AIP (CL062) neurons and major downstream targets. Dendrogram shows the hierarchical clustering of AIP pairs based on downstream connectivity. (B) A circuit diagram of the connectivity between the two AIP (CL062) subtypes and their major downstream targets. Pie charts represent the ratio of input synapses that each downstream target receives from the two AIP subtypes respectively. Only projections from AIP to downstream targets are displayed. (C) A summary table of split-GAL4 drivers screened in this study. Green shades highlight the combinations that were selected for this study based on their specificity and expression strength.

**Figure S2.**
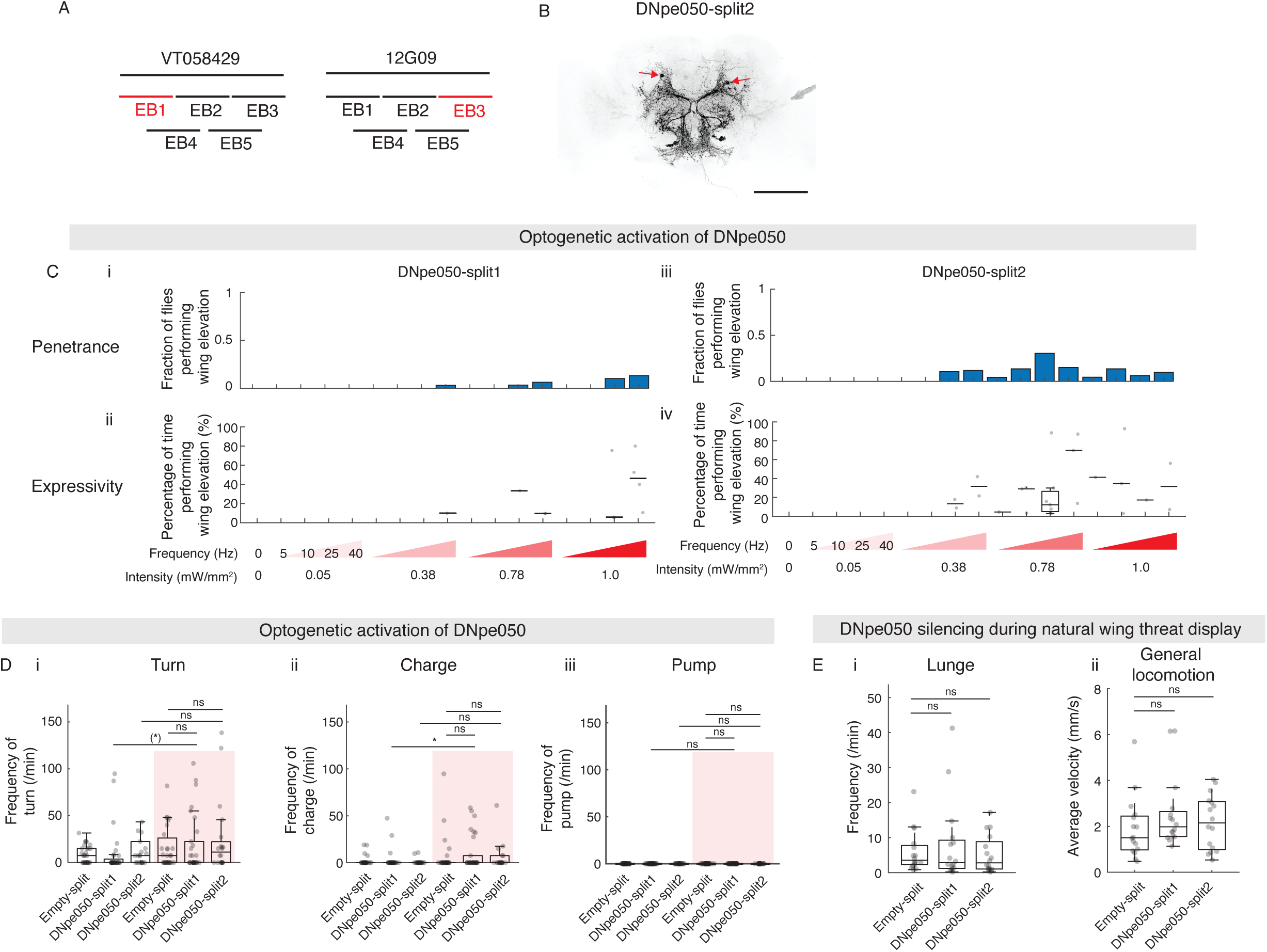
DNpe050 activation does not promote turning, charging, or pumping. (A) A scheme illustrating the enhancer-bashing strategy for identification of sparse and specific DNpe050 drivers. Fragments that retain labeling in DNpe050 are highlighted in red. (B) Expression pattern of DNpe050-split2 (with Otd-FLP). Arrows: cell body locations of DNpe050. Scale bar, 100μm. (C) Fraction of flies performing wing elevation (i) and percentage of stimulation time with wing elevation (ii) during photostimulation of DNpe050-split1 (with Otd-FLP) under various LED frequencies and intensities. (iii and iv) Same quantifications for DNpe050-split2 (with Otd-FLP) activation. Results under 1.0 mW/mm2, 40 Hz activation is duplicated from Figure 2D for the purpose of comparison. (D) DNpe050 (with Otd-FLP) activation does not promote turning (i), charging (ii), or pumping (iii) (E) Silencing of DNpe050 (with Otd-FLP) using Kir2.1 did not impact lunging (i) or average velocity during recording (ii). See also Table S1 and S2.

**Figure S3.**
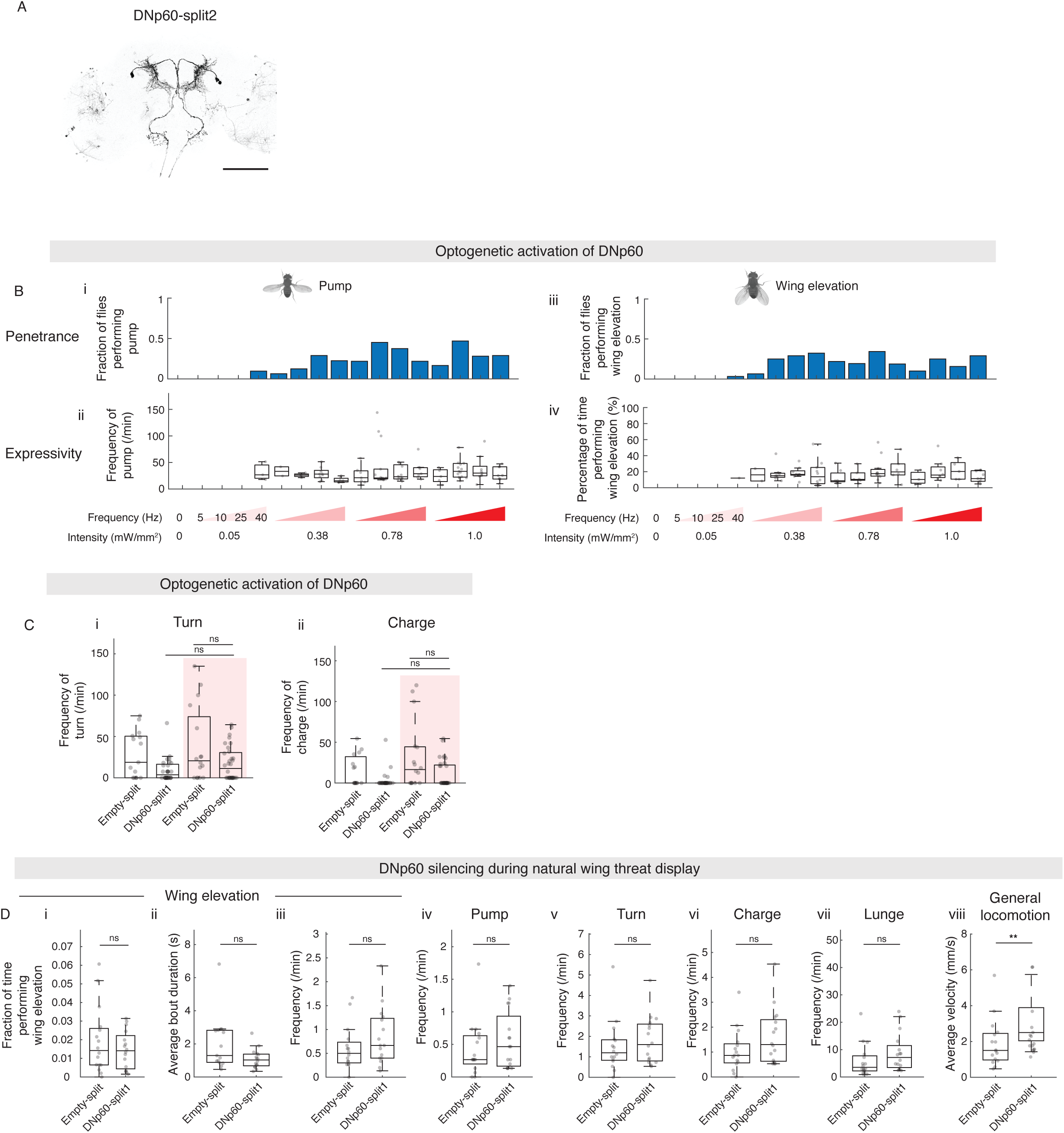
DNp60 does not evoke turning or charging. (A) Expression pattern of DNp60-split2 (with Otd-FLP). Scale bar, 100μm. (B) Fraction of flies performing at least one pump (i), frequency of pump (ii), fraction of flies performing at least one wing elevation bout (iii) and percentage of time performing wing elevation (iv) during photostimulation of DNp60-split1 (with Otd-FLP) under various LED frequencies and intensities. Results under 1.0 mW/mm^2^, 10 Hz are duplicated from Figure 3D and 3E for purposes of comparison. (C) DNp60 (with Otd-FLP) activation does not elicit turning (i) or charging (ii) (D) Behavioral quantification of DNp60 (with Otd-FLP) silencing using Kir2.1: (i) fraction of time performing wing elevation, (ii) average wing elevation bout duration, (iii) frequency of wing elevation, (iv) frequency of pumps, (v) frequency of turns, (vi) frequency of charges, (vii) frequency of lunges, (viii) average velocity during recording. Results of empty-split group are the same as in Figure 2F and S2E. See also Table S1 and S2.

**Figure S4.**
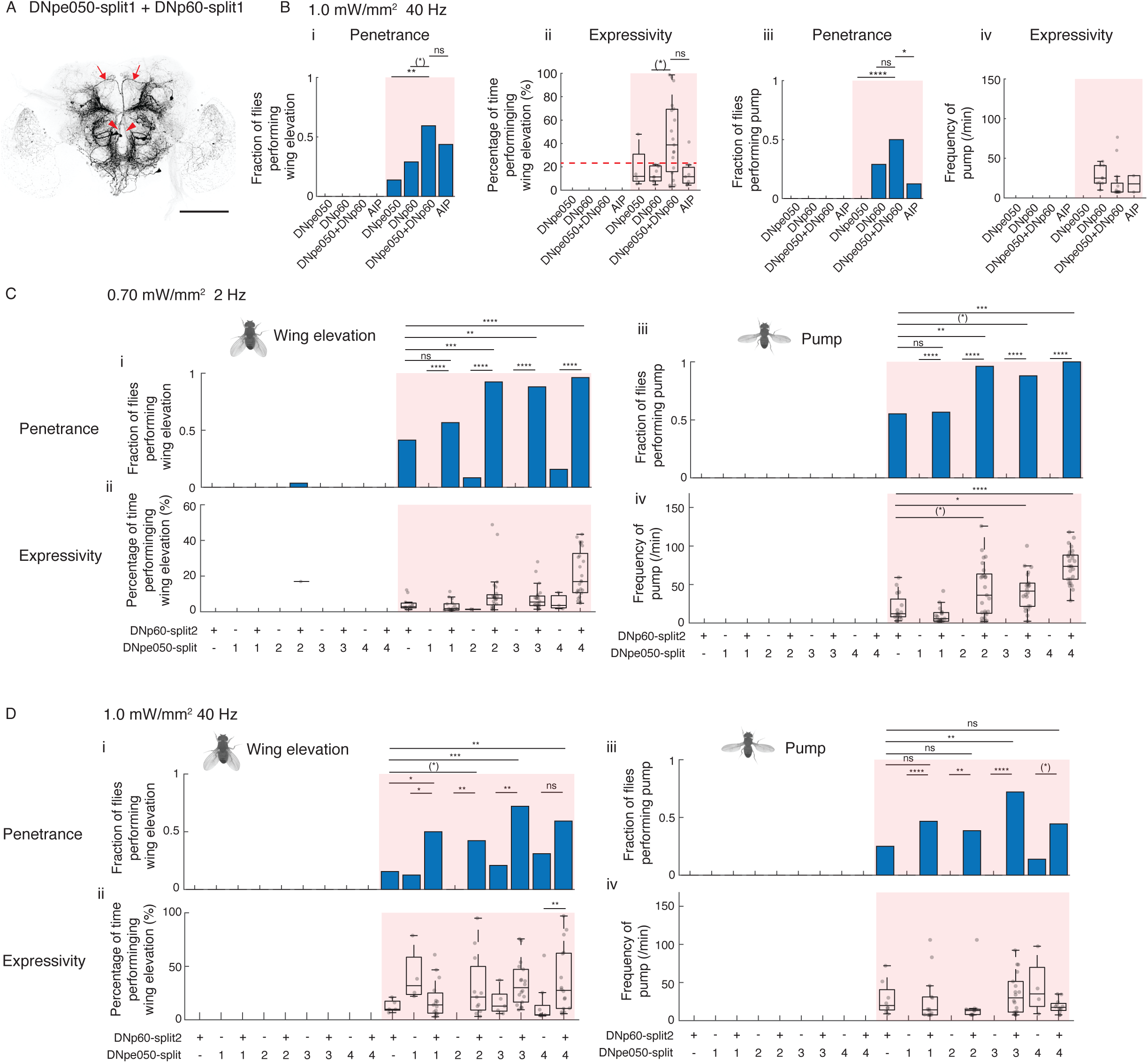
DNpe050 and DNp60 synergistically control wing elevation and pump. (A) Expression pattern of DNpe050-split1 + DNp60-split1 (with Otd-FLP). Scale bar, 100μm. Arrows: descending fibers of DNp60. Arrow heads: descending fibers of DNpe050. (B) Fraction of flies performing at least one wing elevation bout (i), percentage of time performing wing elevation (ii), fraction of flies performing at least one pump (iii), and frequency of pump (iv) during photostimulation of DNpe050 and DNp60 combined (with Otd-FLP, 1.0 mW/mm^2^, 40 Hz, 10 ms pulse width). Results of DNp60 activation are duplicated from Figure S3B for purposes of comparison. (C) Fraction of flies performing at least one wing elevation bout (i), percentage of time performing wing elevation (ii), fraction of flies performing at least one pump (iii), and frequency of pump (iv) during photostimulation of DNp60-split2 combined with DNpe050-split drivers (with Otd-FLP, 0.70 mW/mm^2^, 2 Hz, 30 ms pulse width). (D) Same as (C) but under LED parameters: 1.0 mW/mm^2^, 40 Hz, 10 ms pulse width. See also Table S1 and S2.

**Figure S5.**
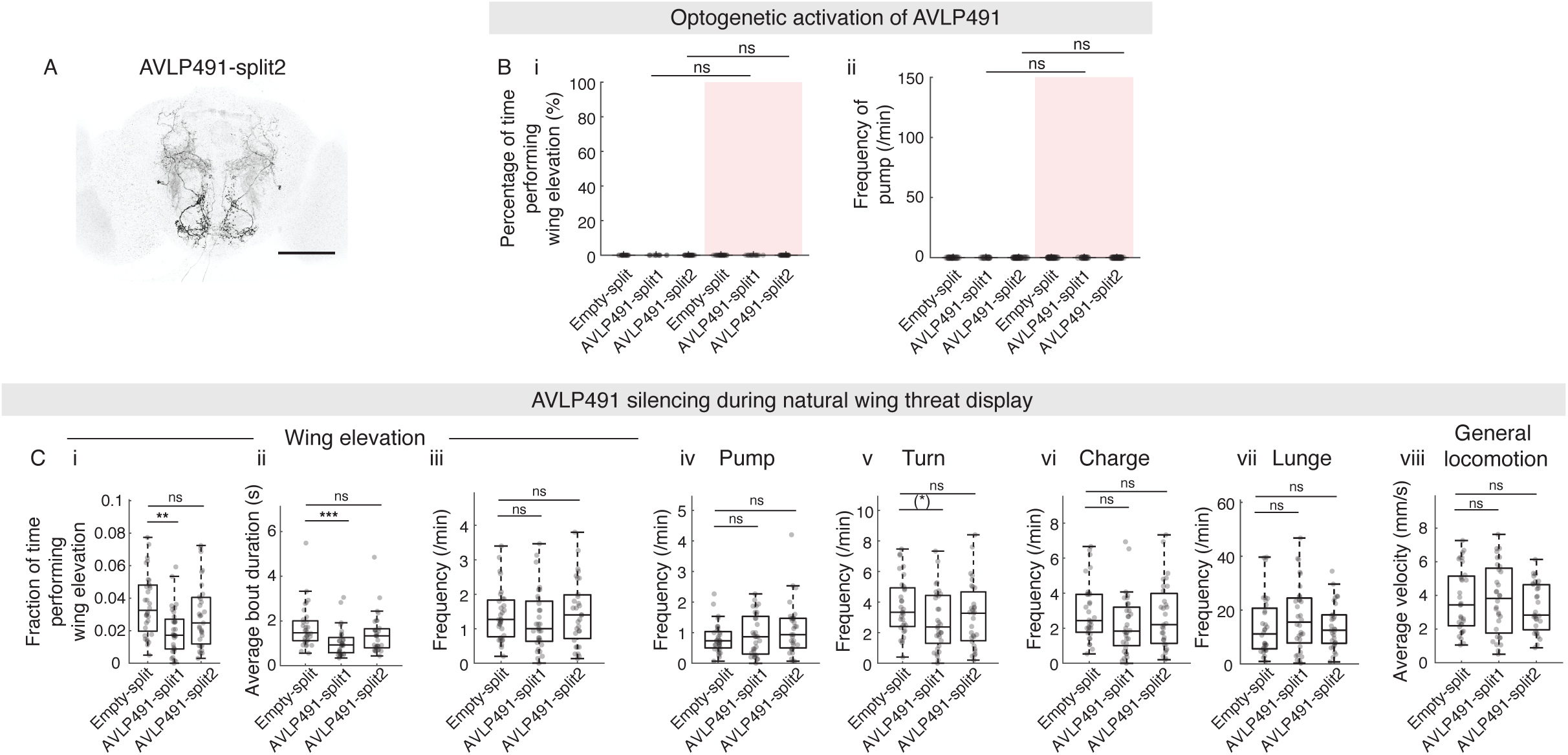
AVLP491 does not induce wing elevation or pump. (A) Expression pattern of AVLP491-split2 (with Otd-FLP). Scale bar, 100μm. (B) Percentage of time flies performing wing elevation (i) and frequency of pump (ii) during AVLP491 (with Otd-FLP) activation. (C) Behavioral quantification of AVLP491 (with Otd-FLP) silencing using Kir2.1: (i) fraction of time performing wing elevation, (ii) average wing elevation bout duration, (iii) frequency of wing elevation, (iv) frequency of pumps, (v) frequency of turns, (vi) frequency of charges, (vii) frequency of lunges, (viii) average velocity during recording. See also Table S1 and S2.

**Figure S6.**
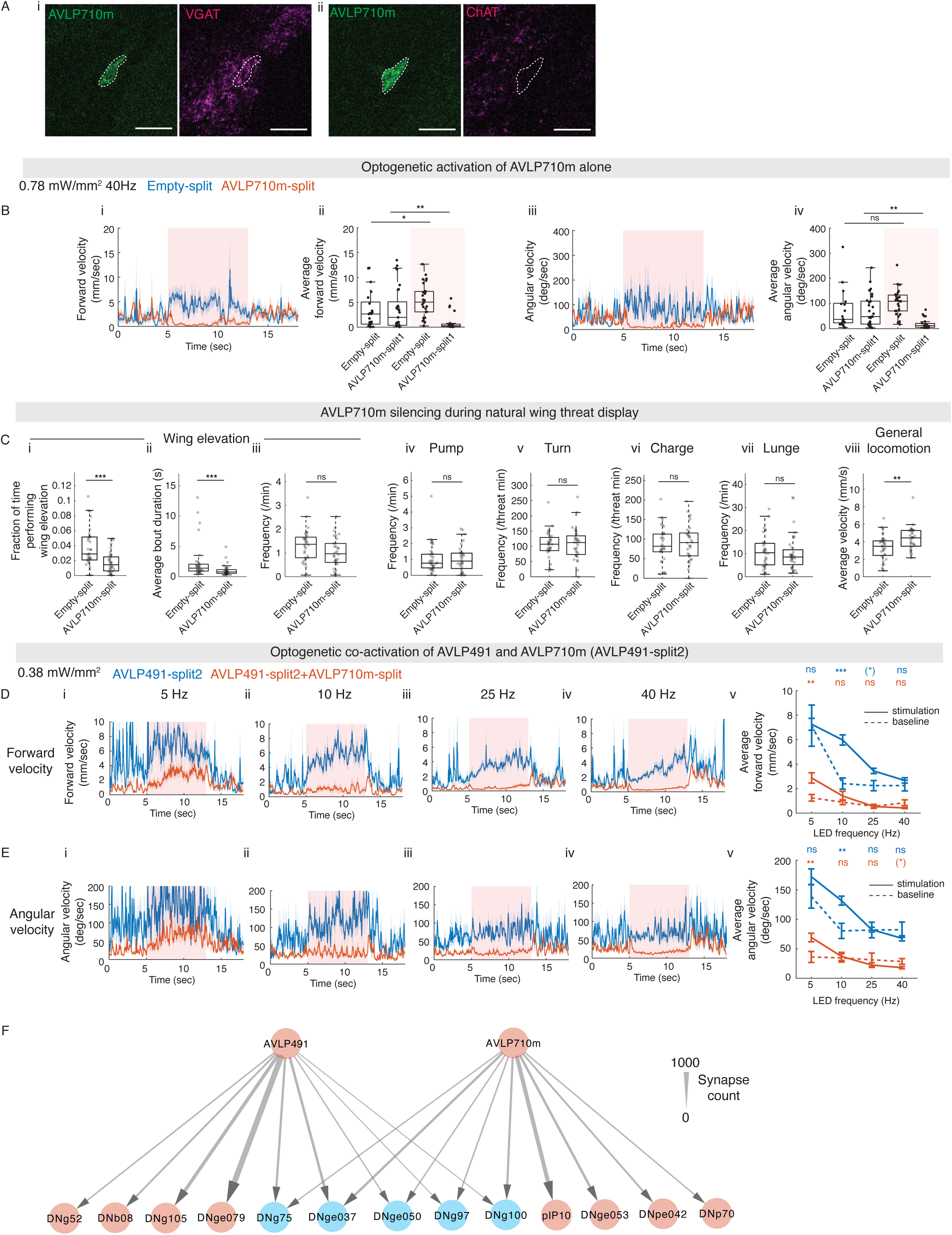
AVLP710m is GABAergic and projects to overlapping set of DNs with AVLP491. (A) Fluorescence *in situ* hybridization for VGAT (i) and ChAT (ii) in AVLP710m. Dashed lines indicate the cell body locations of AVLP710m. Scale bar, 10μm. (B) (i) Forward velocities before, during, and after optogenetic activation of AVLP710m (with Otd-FLP). (ii) Average forward velocities before and during photostimulation. (iii-iv) same as (i-ii) but for angular velocities. Results for empty-split activation are the same as in Figure 5Diii and 5Div. (C) Behavioral quantifications of AVLP710m (with Otd-FLP) silencing using Kir2.1: (i) fraction of time performing wing elevation, (ii) average wing elevation bout duration, (iii) frequency of wing elevation, (iv) frequency of pumps, (v) frequency of turns during wing threat, (vi) frequency of charges during wing threat, (vii) frequency of lunges, (viii) average velocity during recording. (D) Forward velocities before, during, and after optogenetic activation at 5 Hz (i), 10 Hz (ii), 25 Hz (iii) and 40 Hz (iv). (v) Average forward velocities before (dashed line) and during (solid line) photostimulation. Statistic comparisons are performed within genotypes between before (dashed line) and during (solid line) photostimulation. Lines and error bars denote mean ± SEM. (E) same as (D) but for angular velocities. (F) A circuit diagram highlighting the top downstream DN targets of AVLP491 and AVLP710m. Blue circles highlight the common downstream targets. See also Table S1 and S2.

**Figure S7.**
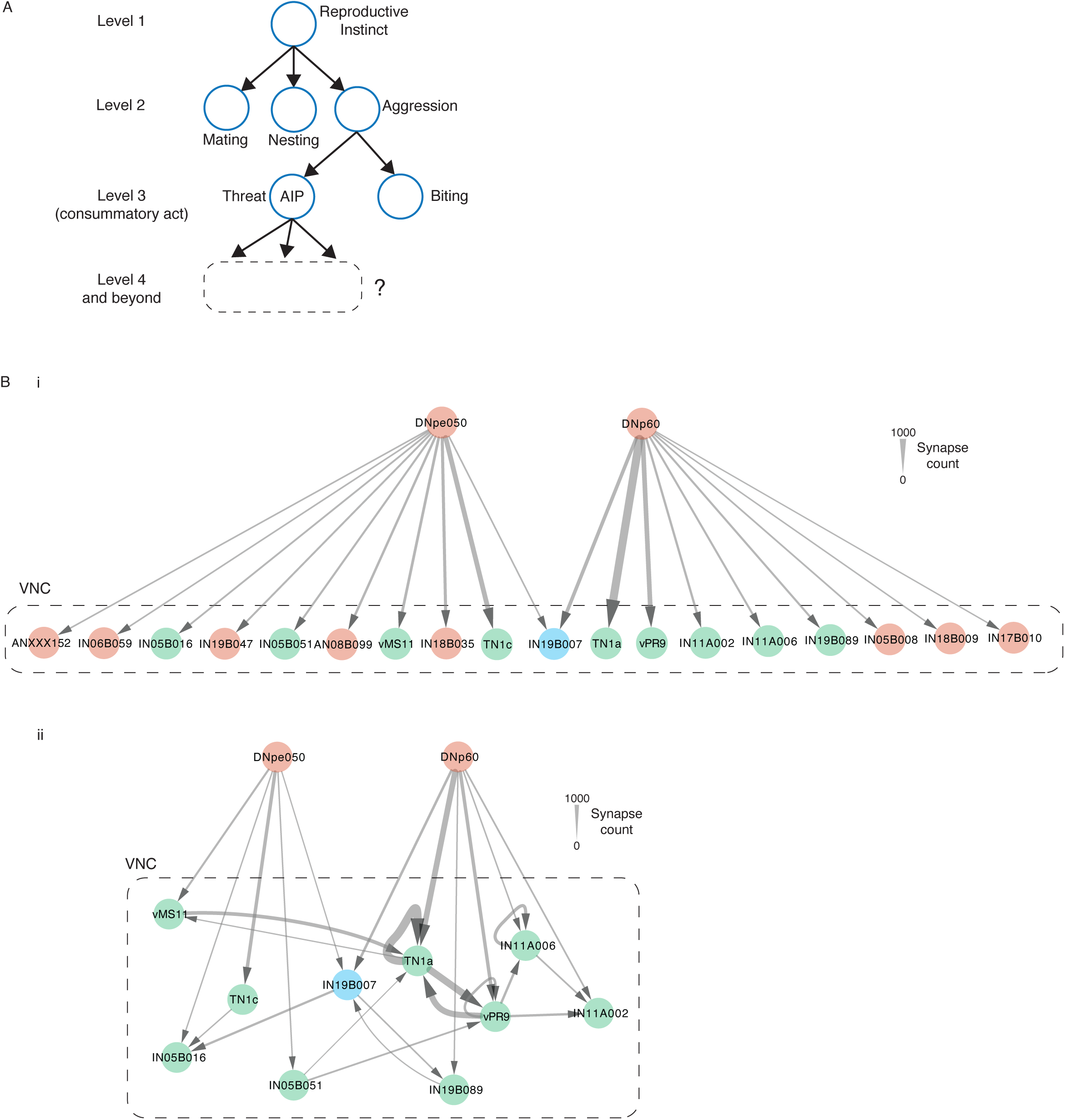
VNC circuit diagram of DNpe050 and DNp60. (A) An illustration of Tinbergen hierarchical behavior control system. (B) Circuit diagrams of DNpe050 and DNp60 downstream VNC targets in Male CNS Connectome. (i) Top downstream VNC targets of DNpe050 and DNp60. Only direct connectivity from DNpe050/DNp60 to downstream targets is displayed. Synapse count cut off: 150 synapses. (ii) A circuit diagram of interconnectivity among DNpe050/DNp60 direct VNC downstream targets. Blue highlights the common downstream target. Green highlights the same neurons displayed in (i) and (ii).

## ACKNOWLEDGMENTS

We thank V. Chiu, C. Schretter, Y. Ouadah, L. Salay, Y. Li, T. Itakura, and Z. Zhong for comments on the manuscript; I. Ros, M. Dickinson, N. Shigehiro, H. Cheong, G. Card, S. Bidaye, B. Dickson, the Janelia Fly Bank, and the Bloomington *Drosophila* Stock Center for fly stocks; Janelia FlyEM Project Team and Cambridge Connectomics Group for sharing information prior to publication; J. Goldammer and G. Rubin for providing the SS33004 line. G. Rubin for advice on connectomics and split-GAL4 generation; S. Berg, S. Takemura and T. Yang for advice on connectomics; M. Mandic for performing preliminary experiments; D. Wagenaar for crafting behavioral chambers; Y. Huang for assisting with plasmid preparation; A. Sanchez for fly maintenance. This work was supported by an NIH grant (R37DA031389) to D.J.A and a Josephine de Karman Fellowship to S.C. D.J.A. is an investigator of the Howard Hughes Medical Institute.

## AUTHOR CONTRIBUTIONS

Conceptualization, S.C and D.J.A.; investigation, S.C.; writing, S.C. and D.J.A.; resources, S.C.; funding acquisition, D.J.A.

## DECLARATION OF INTERESTS

The authors declare no competing interests.

## DECLARATION OF GENERATIVE AI AND AI-ASSISTED TECHNOLOGIES

During the preparation of this work, the authors used ChatGPT (OpenAI) in order to correct grammar mistakes and improve fluency of the manuscript. After using this tool, the authors reviewed and edited the content as needed and take full responsibility for the content of the publication.

## EXPERIMENTAL MODEL AND STUDY PARTICIPANT DETAILS

### Fly Strains

A detailed description of all genotypes used in each experiment is provided in Table S1, and the source of each strain is listed in the Key Resources Table. SS33004 (DNp60-split2), 10xUAS-eGFPKir2.1 (attP2), 20xUAS->myrTopHat2->Chrimson::tdT3.1 (VK5), 20xUAS-Chrimson:tdT(Su(Hw)attP5), 13xLexAop2-Chrimson::tdT3.1 (Su(Hw)attP5), 13xLexAop2-IVS-Syn21-CsChrimson::tdT3.1 (VK5) were kindly shared by the Gerald Rubin laboratory (HHMI Janelia Research Campus) and Barret Pfeiffer. SS33004 (DNp60-split2) was kindly shared by Jens Goldammer and Gerald Rubin. Confocal images of the SS33004 line and other information are available from the Omnibus Broad database^79^. The following strains were obtained from the Bloomington *Drosophila* Stock Center (Indiana University): 10xUAS-IVS-myr::tdTomato3.1 (attP2), 20xUAS-IVS-jGCaMP8m (VK5), 20xUAS-IVS-jGCaMP7b(VK5), VT058429-AD (attP40), 12G09-DBD (attP2), VT061518-DBD (attP2), VT045669-DBD (attP2), 54A01-AD (attP40), VT056582-DBD (attP2), VT016281-DBD (attP2), VT008974-AD (attP40), VT025952-DBD (attP2), 46A02-AD (attP40), 85F12-AD (attP40), VT032899-DBB (attP2), Empty-split (BDP-AD (attP40); BDP-DBD (attP2)).

### Rearing Conditions

Flies were maintained at 25°C and 50% relative humidity on a 12hr:12hr light-dark cycle. Crosses were established with 10-14 virgin females and 3-5 males and transferred to fresh vials every 3-4 days. Experimental flies were collected on day 0-3 post eclosion.

For group-housed condition (used in optogenetic activation and behavioral epistasis experiments), flies were kept at a density of 15-25 individuals per vial. For single-housed condition (used in silencing experiments), flies were reared individually. All experiments were performed 6-8 days after collection.

Flies were reared on standard molasses-based medium. For optogenetic activation and behavioral epistasis experiments, flies were reared in constant darkness on food supplemented with 0.4mM all-trans-retinal after collection and transferred to fresh all-trans-retinal food 1-2 days before testing.

### Construction of Transgenic Animals

The following fly strains were generated in this study: VT058429EB1-AD, VT058429EB2-AD, VT058429EB3-AD, VT058429EB4-AD, VT058429EB5-AD, 12G09EB1-DBD, 12G09EB2-DBD, 12G09EB3-DBD, 12G09EB4-DBD, 12G09EB5-DBD. DNA fragments used for enhancer-bashing were derived from the original VT058429^34,80^ and 12G09^33,81,82^ enhancer sequences and were synthesized by Integrated DNA Technologies, Inc. Cloning procedures followed previous reports^81,82^. Briefly, DNA fragments were inserted into pBPp65ADZpUw (Addgene #26234) and pBPZpGAL4DBD (Addgene #26233) vectors using Gateway cloning and the LR recombination reaction. Germline transformation was performed by BestGene, Inc..

## METHOD

### Immunohistochemistry and Confocal Imaging

Brains or ventral nerve cords were dissected in ice-cold phosphate-buffered saline (PBS) and fixed in 4% paraformaldehyde for 20 min at room temperature. Samples were washed three times in 0.5% PBST (PBS containing 0.5% Triton X-100), for 10 min each at room temperature, then permeabilized in 2% PBST for 30 min at room temperature. Blocking was performed in 5% normal goat serum (NGS) prepared in 0.5% PBST for 1 h at room temperature or overnight at 4°C. Tissues were incubated with primary antibody for at least 48 hours at 4°C (anti-GFP 1:1000, anti-DsRed 1:1000, nc82 1:50), followed by three 10-min washes in 0.5% PBST. Secondary antibodies (1:1000) were applied for 40-48 hours at 4°C, and samples were again washed three times in 0.5% PBST before mounting.

Samples were mounted in Vectashield (Vector Laboratories, Inc.) and imaged on an Olympus Fluoview FV3000 confocal microscope under a 30X objective. Image stacks were acquired at optimal z-step intervals for subsequent analysis. Brain images displayed were maximum intensity projection (MIP) generated in Fiji. Images in Figure 2B and 6B were registered to JRC2018U^83^ using the Fiji CMTK registration GUI^84^ before MIP generation.

### HCR fluorescence In Situ Hybridization

For hybridization chain reaction (HCR) *in situ* hybridization, fly heads were pre-fixed in 4% PFA on ice and dissected in cold PBS. Brains were fixed for 2 hours at 4°C, and HCR v3.0 was performed as previously reported^39^ (also see https://www.moleculartechnologies.org/supp/HCRv3_protocol_generic_solution.pdf), using RNase-free reagents throughout. Following the HCR procedure, brains were post-fixed in 4% PFA for 20 minutes at room temperature, then processed for immunostaining and confocal imaging as described above to label specific cell types.

### Behavioral Assays

All behavioral experiments were conducted at 25°C and 50% relative humidity during zeitgeber time (ZT) 0-4h. Behavioral chambers were illuminated from below with 850mm infrared backlights (SOBL-200x150-850, Smart Vision Lights). Videos were recorded from above at 30 frames/s using Blackfly S USB3 cameras (BFS-U3-13Y3M-C) equipped with long-pass infrared filters (LP780, Midwest Optical Systems). Two types of chambers were used. For experiments in Figure 7B, 7C, 7E, 7F, and 7G, a Heisenberg chamber^16,85^ measuring 47mm (length) x 38mm (width) x 60mm (height) was used; the central 24 mm diameter region was coated with apple juice agar (2.25% (w/v) agarose and 2.5% (w/v) sucrose in apple juice). For most of the experiments, an 8-well chamber was used, consisting of 12mm high x 16 mm diameter acrylic cylinders^86^. The floor of the chamber was uniformly coated with apple juice agar.

For optogenetic activation and behavioral epistasis assays, when using 8-well chambers, one male fly was loaded per well with a mouth aspirator. Optogenetic stimulation was delivered using a custom LED setup as described previously^86^, with a 655nm LED (SR-05-D000, Luxeon Star LED) positioned ∼8 cm above each well at a 24° angle. The stimulation protocol began with a 15 s baseline, followed by sixteen 8 s light periods, each separated by a 15 s interval. For optogenetic activation experiments, stimulation parameters consisted of all combinations of 5, 10, 25, and 40 Hz with light intensities of 0.05, 0.38, 0.78, 1.0 mW/mm^2^, each light pulse lasting 10 ms, or all combinations of 1, 2, 3, 4 Hz with light intensities of 0.45, 0.7, 0.95, 1.2 mW/mm^2^, each light pulse lasting 30 ms for low frequency stimulation. For behavioral epistasis experiments, stimulation parameters consisted of all combinations of 1, 7, 13, 19, 25, 31, and 37 Hz with intensities of 0.18 and 0.50 mW/mm^2^, each pulse lasting 10 ms. In both cases, the total recording time was 420 s. When using the Heisenberg chamber for optogenetic activation, 7-10 male flies were loaded for each recording. Two arrays of four LEDs each delivered light from below at a 24° angle to ensure uniform illumination across the chamber. The stimulation protocol consisted of a 15 s baseline, 24 s stimulation periods and 15 s intervals, for a total recording time of 420 s with nine stimulation trials. Stimulation parameters included all combinations of 2, 3, and 4 Hz with intensities of 0.43, 0.70, and 0.90 mW/mm^2^, and individual pulse duration was 30 ms.

For Kir2.1 silencing experiments, flies were single housed immediately after collection under. Two males of the same genotype were introduced into each well of the 8-well chamber. Each recording lasted 15 minutes.

In all experiments, the chamber walls were coated with Insect-A-Slip or Fluon (Tar Heel Ants, Raleigh, NC), and the lids were coated with silicon fluid to prevent climbing, and flies were allowed to acclimate for 4 min after being transferred into the chamber before the video recording started.

### Two-photon functional imaging

Two-photon calcium imaging was performed following previously described procedures^87^ with minor modifications. Flies were briefly anesthetized on ice for 20-30 s and transferred to an aluminum plate with a open slit that physically restrained the body. The plate was kept cool with ice throughout preparation. Each fly was then mounted onto a custom holder^88^ using UV-cured glue to secure the head, and the proboscis was glued in place to minimize brain motion. The holder was then filled with physiological saline (108 mM NaCl, 5 mM KCl, 4 mM NaHCO_3_, 1 mM NaH_2_PO_4_, 5 mM trehalose, 10 mM sucrose, 5 mM HEPES, 0.5 mM CaCl_2_, 2 mM MgCl_2_, pH = 7.5). A small piece of head cuticle, along with overlying trachea and adipose tissues, was removed to expose the brain, and the esophagus was severed to further reduce movement artifacts.

Calcium imaging was conducted on a custom-modified Ultima two-photon laser-scanning microscope (Bruker). Cell bodies of interest were scanned at 3 frames/s using a 920 nm laser (Chameleon Ultra II Ti:Sapphire laser, Coherent), and images were collected through a 25x objective (XLPlan N, Olympus) with a numerical aperture of 1.00.

For photostimulation, a 655nm LED was placed at ∼1 cm below the fly. Recording includes a 30 s baseline period, 5 s stimulation periods, each separated by a 55 sec recovery interval. Stimulation parameters consist of all combinations of 5, 10, 20, and 40 Hz at the intensities of 0.3, 0.5, and 0.9 mW/mm^2^. The LED was controlled and synchronized with image acquisition via an Arduino-based interface and custom-written scripts. To minimize leak-through into the imaging channel, the LED source was covered with three layers of Roscolux 26 filters (Rosco).

## QUANTIFICATION AND STATISTICAL ANALYSIS

### Statistics

All statistical analyses were performed in MATLAB. Table S2 provides a complete summary of the sample size (n) for each experiment, the statistical test used, the resulting *p*-values, and the adjusted α significance threshold when Bonferroni’s multiple comparison correction was applied.

### Connectome analysis

Analyses were performed using the male CNS connectome dataset^27^ (https://neuprint.janelia.org/, male-cns:v0.9), accessed through the neuPrint interface^89^ and the natverse package^90^ in R. Connections between neuron pairs with five synapses or fewer were excluded. The remaining connections were used to generate a connectivity table, which was imported into Cytoscape (cytoscape.org) for circuit diagram construction. In these diagrams, neurons of the same type from both hemispheres were grouped into a single node, and synapse counts were summed to represent the connection strength between two cell types. Reconstructed neuron skeleton were registered to the JRC2018U brain template^83^. Neuronbridge^29^ was used to identify candidate hemi-drivers, and Split-GAL4s combinations were subsequently generated for anatomical and functional screening.

For AIP subtype analysis, downstream targets included the neuron types with the highest synaptic connectivity to AIP, as well as neurons identified as functionally important for threat behavior (e.g. DNp60). Paris of AIP neurons were grouped into single types based on morphology, and their synapse counts were summed. Heatmaps displaying synapse counts between neuron types were generated with ComplexHeatmap package^91^ in R. Hierarchical clustering was performed with ComplexHeatmap using the complete-linkage method based on Euclidean distance. AIP1 and AIP2 were selected from the highest branching level of the resulting dendrogram.

### Fly tracking and behavior classification

Behavioral recordings were processed with Caltech FlyTracker^92^ to extract various features from the videos such as the positions and the velocities of the flies.

For optogenetic activation and behavioral epistasis experiments, threat action classification followed previous report^16^ with additional filtering step to improve accuracy in the assays used in this study. Periods during which flies remained vertically on the behavior chamber wall were excluded from analysis, as wall climbing often caused tracking errors. Stable wing elevation was defined as both wings extended between 35° and 65° for more than 0.2 s. Pump was defined as average wing angle exceeding 65°. Turning and charging was classified by identifying bouts where angular velocity exceeded 75°/s and forward velocity exceeded 6 mm/s, respectively. To specifically identify discrete events, bouts occurring during periods of sustained locomotion (>20°/s angular velocity or >2mm/s forward velocity for more than 0.4 s) were removed, retaining only discrete and brief events. For Figure 7G, the forward velocity threshold for charging-like behavior was set to 3 mm/s, as coactivation did not generate as high forward velocities characteristic of AIP activation or natural wing threat. Behavioral frequencies were calculated as the number of bouts divided by the total time during photostimulation when flies were not on the wall. 3, 5 or 8 seconds prior to the stimulation onset were used as baseline. For behavioral epistasis experiments, stimulation trials where genetic controls displayed the highest levels of threat behavior were used for quantification. In optogenetic activation experiments, false-positive wing elevation and pump events were manually removed.

For Kir2.1 silencing experiments examining natural wing threat, a JAABA classifier was used to detect bouts of natural wing threat, and lunging was identified with a previously published JAABA classifier^22^. Misidentified bouts resulting from wall climbing were excluded. Each wing threat bout was counted as a single wing elevation bout. All classified wing threat bouts were then manually proofread. For identifying pump, turning, and charging, a 0.5 s window before and after each wing elevation bout was included for analysis, following the convention of our previous study^16^, and the same criteria and filtering were applied as described above. Action frequency was calculated as the number of bouts divided by the total recording duration. When there were significant differences in fraction of time performing wing elevation, frequency of turns and charges were normalized to the length of analysis time window (wing elevation period plus 0.5 s before and after).

### Pearson Correlation Coefficient

Periods when flies were positioned vertically on the chamber wall were excluded from analysis. To remove artifacts from extreme movements such as jumping or flying, frames in which angular velocity exceeded 1,000°/s or forward velocity exceeded 100 mm/s were also excluded. If two flies share less than 10% overlap in post-filtered recording time, the Pearson correlation coefficient was not computed and was recorded as *Not a Number* (NaN). Cumulative distribution plots were generated using all non-NAN data points. Statistical significance between two genotypes was assessed via a random-labeling bootstrap implemented in a custom MATLAB script. In each iteration, genotype labels were shuffled across all individuals, and the difference in the medians of the Pearson correlation coefficients between two groups was calculated. This process was repeated for 10,000 times. The *p*-value was computed as the proportion of the iterations in which the simulated difference exceeded the observed difference.

### Two-photon imaging data analysis

After image acquisition, a circular region of interest (ROI) was manually drawn in Fiji to encompass the cell body. Pixel intensity within the ROI was measured across the entire recording session using a custom written MATLAB script, yielding the raw fluorescence trace. The first 20 seconds of each recording were used to establish the baseline fluorescence (F_0_). Changes in fluorescence were then expressed as dF/F, calculated relative to this baseline.

## Notes

### Competing Interest Statement

The authors have declared no competing interest.

